# Regulatory network analysis of Paneth cell and goblet cell enriched gut organoids using transcriptomics approaches

**DOI:** 10.1101/575845

**Authors:** A Treveil, P Sudhakar, Z J Matthews, T Wrzesinski, E J Jones, J Brooks, M Olbei, I Hautefort, L J Hall, S R Carding, U Mayer, P P Powell, T Wileman, F Di Palma, W Haerty, T Korcsmáros

## Abstract

The epithelial lining of the small intestine consists of multiple cell types, including Paneth cells and goblet cells, that work in cohort to maintain gut health. 3D *in vitro* cultures of human primary epithelial cells, called organoids, have become a key model to study the functions of Paneth cells and goblet cells in normal and diseased conditions. Advances in these models include the ability to skew differentiation to particular lineages, providing a useful tool to study cell type specific function/dysfunction in the context of the epithelium. Here, we use comprehensive profiling of mRNA, microRNA and long non-coding RNA expression to confirm that Paneth cell and goblet cell enrichment of murine small intestinal organoids (enteroids) establishes a physiologically accurate model. We employ network analysis to infer the regulatory landscape altered by skewing differentiation, and using knowledge of cell type specific markers, we predict key regulators of cell type specific functions: Cebpa, Jun, Nr1d1 and Rxra specific to Paneth cells, Gfi1b and Myc specific for goblet cells and Ets1, Nr3c1 and Vdr shared between them. Links identified between these regulators and cellular phenotypes of inflammatory bowel disease (IBD) suggest that global regulatory rewiring during or after differentiation of Paneth cells and goblet cells could contribute to IBD aetiology. Future application of cell type enriched enteroids combined with the presented computational workflow can be used to disentangle multifactorial mechanisms of these cell types and propose regulators whose pharmacological targeting could be advantageous in treating IBD patients with Crohn’s disease or ulcerative colitis.

**Table of contents:** We demonstrate the application of network biology techniques to increase understanding of intestinal dysbiosis through studying transcriptomics data from Paneth and goblet cell enriched enteroids.

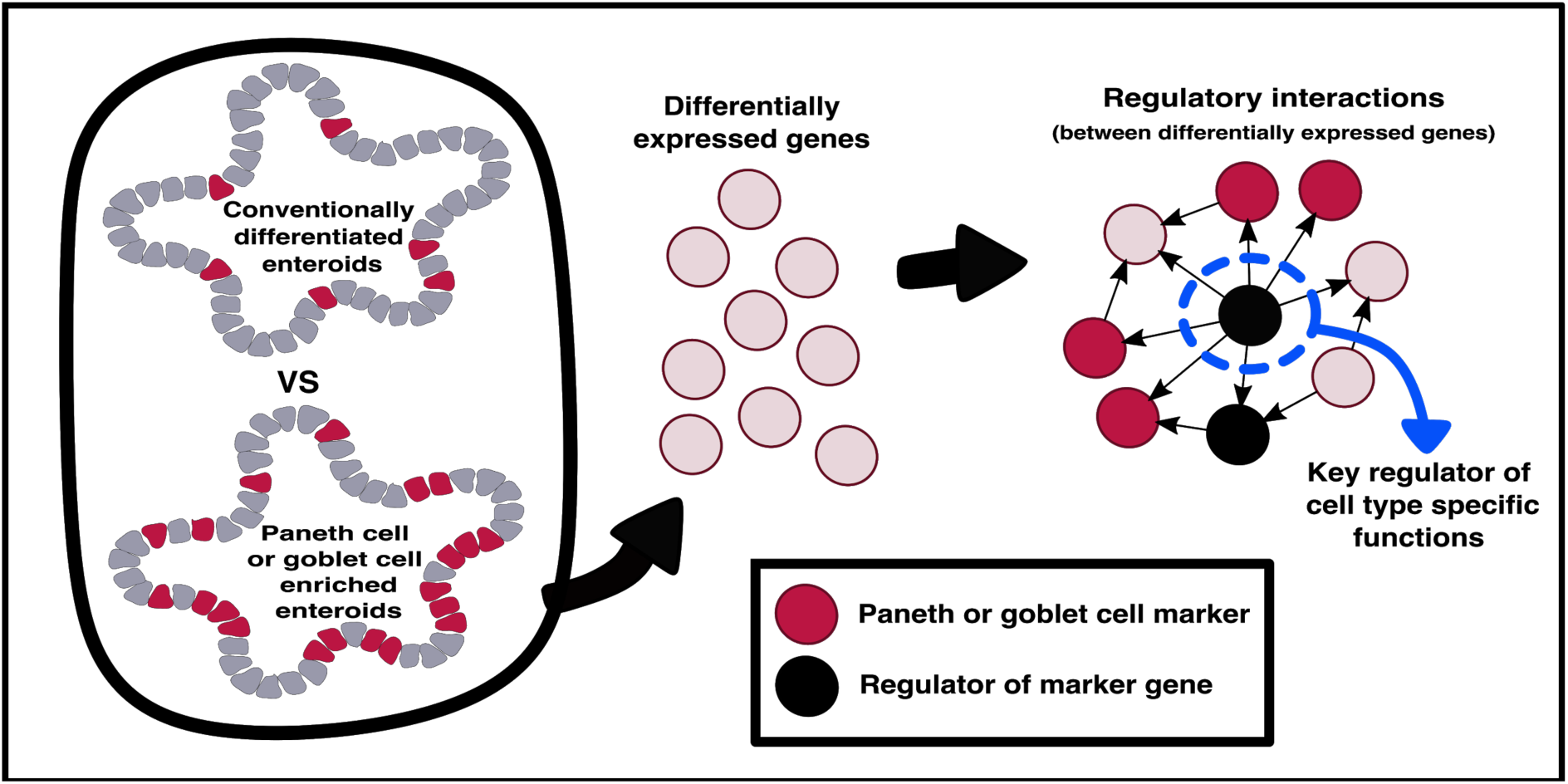

## Introduction

Gut barrier integrity is critically important for efficient nutrient absorption and maintenance of intestinal homeostasis (1) and is maintained by the combined action of the various cell types lining the intestinal epithelium (2). These intestinal epithelial cells serve to mediate signals between the gut microbiota and the host innate/adaptive immune systems (3,4). Disruption of epithelial homeostasis along with dysregulated immune responses are some of the underlying reasons behind the development of inflammatory bowel disease (IBD) such as Crohn’s disease (CD) and ulcerative colitis (UC) (5). Therefore, a greater insight into the functions of intestinal cells will further our understanding of the aetiology of inflammatory gut conditions.

To date, various cell types have been identified in the intestinal epithelium based on specific functional and gene expression signatures. Paneth cells residing in the small intestinal crypts of Lieberkühn help to maintain the balance of the gut microbiota by secreting anti-microbial peptides, cytokines and other trophic factors (6). Located further up the intestinal crypts, goblet cells secrete mucin, which aggregates to form the mucus layer, which acts as a chemical and physical barrier between the intestinal lumen and the epithelial lining (7). Both of these cell populations have documented roles in gut-related diseases (8,9). Dysfunctional Paneth cells with reduced secretion of anti-microbial peptides have been shown to contribute to the pathogenesis of CD (10), while reduction in goblet cell numbers and defective goblet cell function has been associated with UC in humans (11).

Recent studies have employed single cell transcriptomics sequencing of tissue samples to characterise the proportion and signatures of different epithelial cell types in the intestines of healthy and IBD patients (12–14). However, to provide deeper insights into the role of specific cell populations (such as Paneth cells and goblet cells) in IBD, *in vitro* models are required for in depth testing and manipulation. Such models can be used to study specific mechanisms of action, host-microbe interactions, intercellular communication, patient specific therapeutic responses and to develop new diagnostic approaches. Due to ease of manipulation, observation and analysis, organoid models, including small intestinal models (enteroids), are increasingly used in the IBD field (15–17). Enteroid cell culture systems employ growth factors to expand and differentiate Lgr5+ stem cells into spherical models of the small intestinal epithelia which recapitulate features of the *in vivo* intestinal tissue (18–20). It has been shown that these enteroids contain all the major cell types of the intestinal epithelium, and exhibit normal *in vivo* functions (21). These models, generated using intestinal tissue from mice, from human patient biopsies or from induced pluripotent stem cells (iPSCs), have proven particularly valuable for the study of complex diseases which lack other realistic models and exhibit large patient variability, such as IBD (22,23). Small molecule treatments have been developed that skew the differentiation of enteroids towards Paneth cell or goblet cell lineages, improving representation of these cells within the enteroid cell population (24,25). Specifically, differentiation can be directed towards the Paneth cell lineage through the addition of DAPT, which inhibits notch signaling, and CHIR99021, which inhibits GSK3β-mediated β-catenin degradation. Enteroid cultures enriched in goblet cells can be generated through the addition of DAPT and IWP-2, an inhibitor of Wnt signaling (25). Whilst these methods do not present single cell type resolution, they provide useful tools to study Paneth cell and goblet cell populations in the context of the other major epithelial cell types (26). A recent study by Mead et al. found that Paneth cells from enriched enteroids more closely represent their *in vivo* counterparts than those isolated from conventionally differentiated enteroids, based on transcriptomics, proteomics and morphologic data (27). Furthermore, we have shown that enteroids enriched for Paneth cells and goblet cells recapitulate *in vivo* characteristics on the proteomics level (28,29) and that they are a useful tool for the investigation of health and disease related processes in specific intestinal cell types (29).

Nevertheless, the effect of Paneth cell and goblet cell enrichment of enteroids on key regulatory landscapes has not been extensively characterised. In this study, we comprehensively profiled mRNAs, miRNAs and lncRNAs from mouse derived, 3D conventionally differentiated enteroids (control), Paneth cell enriched enteroids (PCeE) and goblet cell enriched enteroids (GCeE) to determine the extent to which these enteroids display increased Paneth cell and goblet cell signatures. We applied a systems-level analysis of regulatory interactions within the PCeEs and GCeEs to further characterise the effect of cell type enrichment and to predict key molecular regulators involved with Paneth cell and goblet cell specific functions. This analysis was carried out using interactions networks, which are a primary method to collate, visualise and analyse biological systems. These networks are a type of systems biology data representation, which aids the interpretation of -omics read-outs by contextualising genes/molecules of interest and identifying relevant signalling and regulatory pathways (30,31). In the presented analysis, the nodes of the interaction networks represent genes/molecules of interest from the transcriptomics data and the edges represent regulatory connections (molecular interactions) between the nodes inferred from databases. Studying regulatory interactions using interaction networks has been proved useful to uncover how cells respond to changing environments at a transcriptional level, to prioritise drug targets and to investigate the downstream effects of gene mutations and knockouts (32–34).

We used network approaches to interpret the PCeE and GCeE transcriptomic data by integrating directed regulatory connections from resources containing transcriptional and post-transcriptional interactions. This integrative strategy led us to define regulatory network landscapes altered by Paneth cell and goblet cell enrichment of enteroids, termed as PCeE network and GCeE network, respectively (Figure 1). By incorporating known Paneth cell and goblet cell markers, we used these networks to predict master regulators of Paneth cell and goblet cell differentiation and/or maintenance in the enriched enteroids. Furthermore, we highlighted varying downstream actions of shared regulators between the cell types. This phenomenon, called regulatory rewiring, highlights the importance of changes in regulatory connections in the function and differentiation of specific cell types. Taken together, we show that cell type enriched enteroids combined with the presented network biology workflow have potential for application to the study of epithelial dysfunction and mechanisms of action of multifactorial diseases such as IBD.

**Figure 1:**
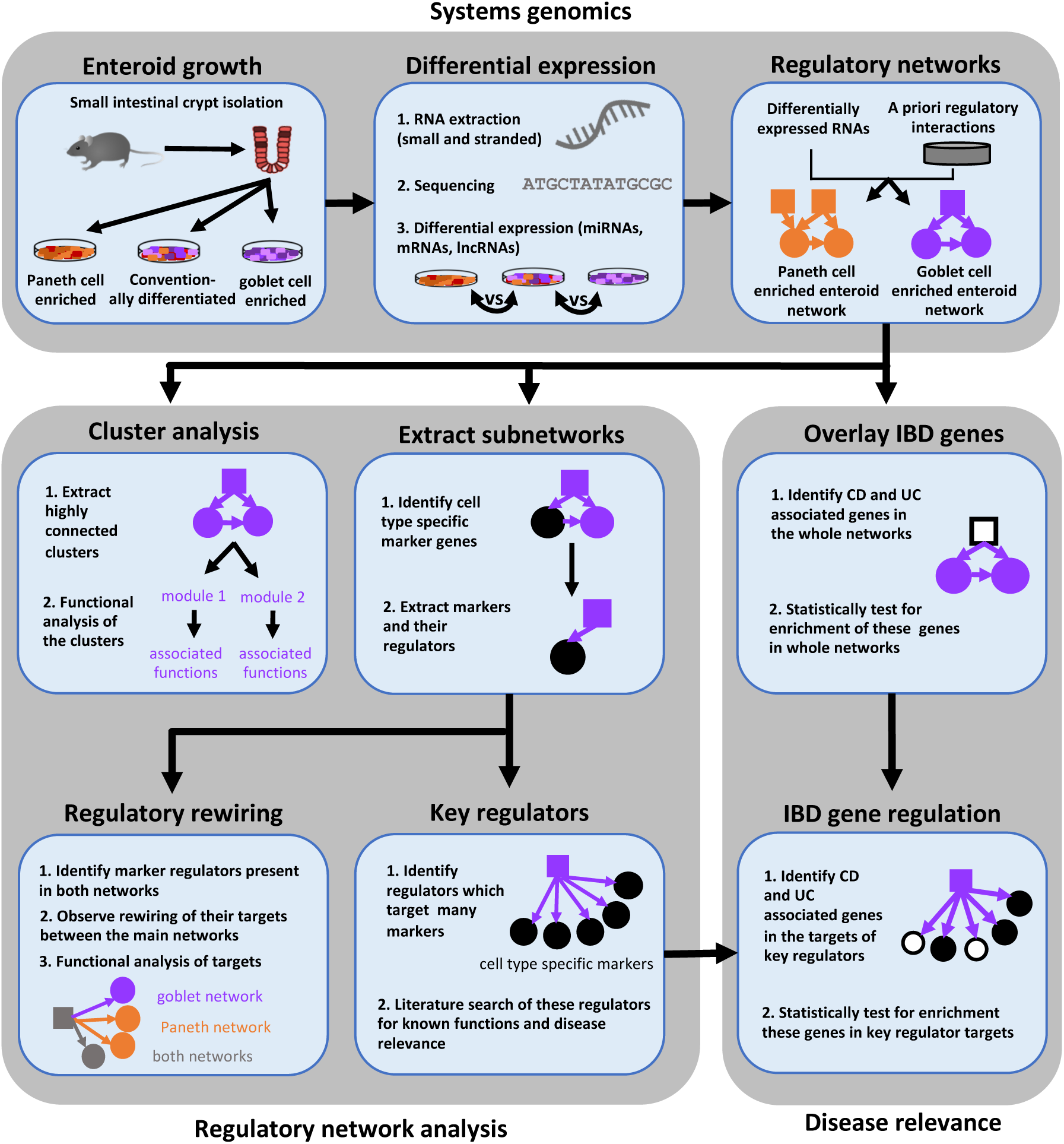
Schematic representation of the workflow used to infer and analyse regulatory network landscapes altered by Paneth cell and goblet cell enrichment of enteroids. PCeE/GCeE network - Paneth cell enriched enteroid / goblet cell enriched enteroid network; TF- transcription factor; lncRNA-long non-coding RNA; miRNA-microRNA; mRNA-messenger RNA; UC - ulcerative colitis; CD - Crohn’s disease.

## Results

### Secretory lineages are over-represented in cell type enriched enteroids compared to conventionally differentiated controls

We generated 3D self-organising enteroid cultures *in vitro* from murine small intestinal crypts (Figure S1) (18–20). In addition to conventionally differentiated enteroids, we generated enteroids enriched for Paneth cells and goblet cells using well-established and published protocols, presented in detail in the Methods (20,25). Bulk transcriptomics data was obtained from each set of enteroids to determine genes with differential expression resulting from enteroid skewing protocols. Differentially expressed genes were calculated by comparing the RNA expression levels (including protein coding genes, lncRNAs and miRNAs) of enteroids enriched for Paneth cells or goblet cells to those of conventionally differentiated enteroids. 4,135 genes were differentially expressed (absolute log2 fold change ≥ 1 and false discovery rate ≤ 0.05) in the PCeE dataset, and 2,889 were differentially expressed in the GCeE dataset (Figure 2A-C, Table S1). The larger number of differentially expressed genes (DEGs) in the PCeE data could be attributed to the highly specialised nature of Paneth cells (35,36). The majority of the DEGs were annotated as protein coding: 79% in the PCeE dataset and 84% in the GCeE dataset. In addition, we identified lncRNAs (PCeE, 11%; GCeE, 9%) and miRNAs (PCeE, 4%; GCeE, 2%) among the DEGs (Figure 2B). Some of these DEGs were identified in both the PCeE and the GCeE datasets, exhibiting the same direction of change compared to the conventionally differentiated enteroid data. In total, 1,363 genes were found upregulated in both the PCeE and the GCeE data, while 442 genes were found downregulated in both datasets (Figure 2C). This result highlights considerable overlap between the results of skewing enteroids towards Paneth cells and goblet cells, and can be explained by the shared differentiation history and secretory function of both Paneth cells and goblet cells.

**Figure 2:**
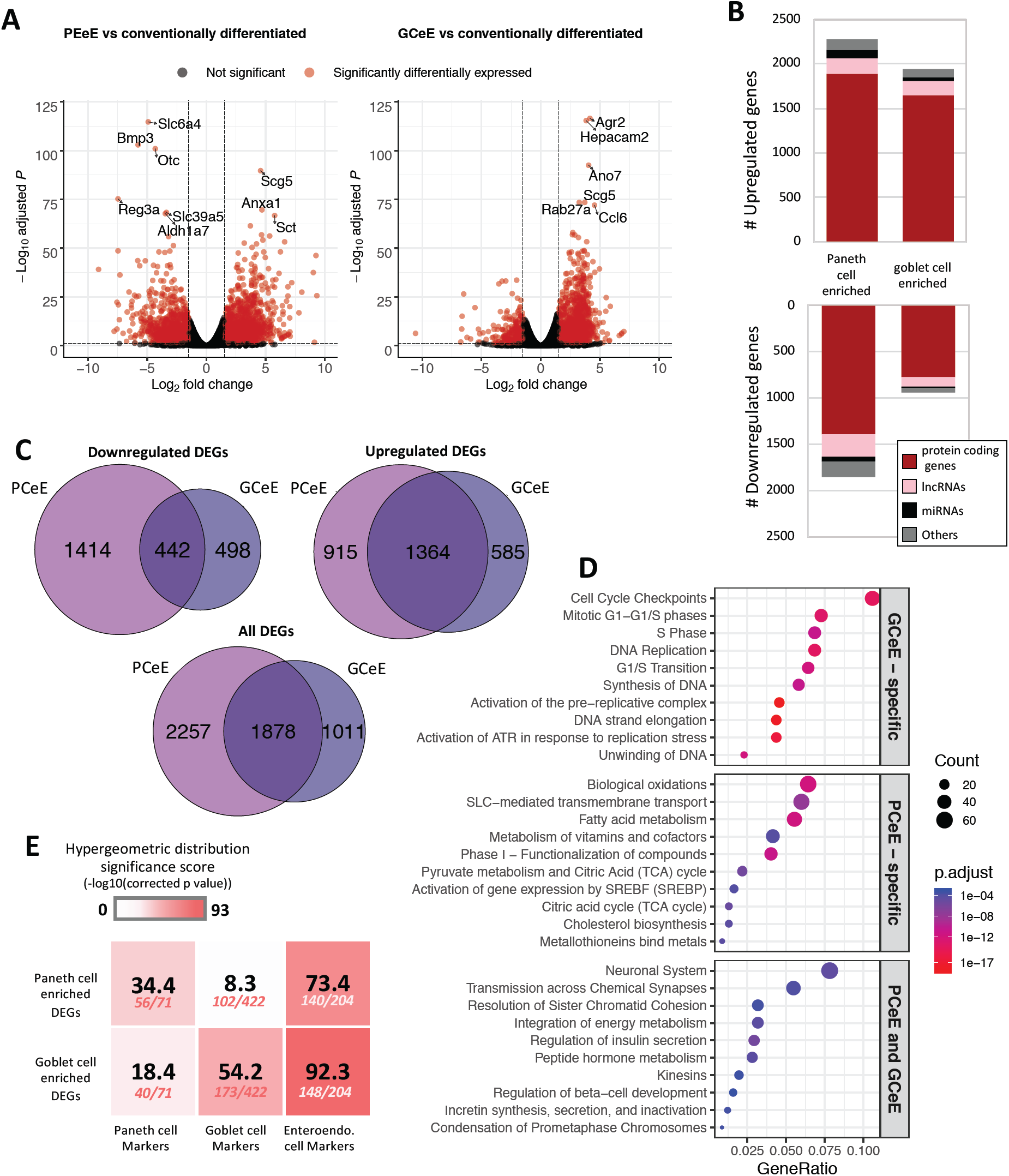
Differentially expressed genes in Paneth cell enriched enteroids (PCeEs) and goblet cell enriched enteroids (GCeEs) (compared to conventionally differentiated enteroids). **A:** Volcano plots showing log2 fold change and adjusted p value for each gene following differential expression analysis of PCeEs (left) and GCeEs (right). Horizontal and vertical lines indicate the differential expression criteria cut offs (q value ≤ 0.05 and log2 fold change ≥|1|). **B:** Number of differentially expressed genes (DEGs). miRNA = microRNA; lncRNA = long non-coding RNA; Genes annotated as ‘other’ include pseudogenes and antisense genes. **C:** Venn diagrams indicating the number of DEGs (passing the cut off criteria). **D:** Top 10 Reactome pathways of the 50 most significant DEGs (by q value). **E:** Enrichment of cell type specific marker genes in the DEG lists. Higher significance scores indicate greater enrichment. Number of markers in DEG list out of the total number of markers shown below significance score. Also see Table S1,4.

Pathway analysis was employed to study functional associations of the DEGs (Figure 2D). The PCeE-specific DEGs were associated with a number of metabolic pathways, including Metabolism of vitamins and cofactors, Pyruvate metabolism and Citric Acid (TCA) cycle and Cholesterol biosynthesis. On the other hand, GCeE-specific DEGs were associated with the cell cycle through pathways such as Cell Cycle Checkpoints, DNA replication and G1/S transition. Pathways associated with the shared DEGs included Transmission across Chemical Synapses, Integration of energy metabolism and a number of pathways linked to hormones. As hormone functions are characteristic of enteroendocrine cells, this analysis suggests that enteroendocrine cells are enriched in both the PCeEs and the GCeEs.

To validate the cell types present in the enteroids, the expression of five previously reported major cell type specific markers were investigated across the enteroids using transcript abundances and RNA differential expression results (Table S2, Figure S2). The control enteroids and the cell type enriched enteroids expressed all five investigated markers: *Lgr5* (stem cells), *ChgA* (enteroendocrine cells), *Muc2* (goblet cells), *Lyz1* (Paneth cells) and *Vil1* (epithelial cells). We observed an upregulation of *Muc2, Lyz1* and *ChgA* and a downregulation of *Lgr5* in PCeEs and GCeEs compared to the control enteroids, confirming the more pronounced differentiated status of the enteroids. In addition, a number of Paneth cell specific antimicrobial peptide genes were differentially expressed in the PCeE dataset, including *Ang4, Reg3γ, Pla2g2a and Defa2* (Table S3). Some of these genes were also differentially expressed in the GCeE dataset but with smaller log fold change values, e.g. *Lyz1* and *Ang4*. Conversely, a number of goblet cell mucin related genes (including *Muc2* and *Tff3*) were differentially expressed in both datasets although all genes exhibited a smaller increase in the PCeEs (Table S2). Therefore, using primary cell type specific markers, antimicrobial peptide genes and mucin-related genes, we show that the enteroids contain all major cell types, and that Paneth cell are most upregulated in the PCeEs, while goblet cells are most upregulated in the GCeEs. We also note that both differentiation methods resulted in increases of other secretory cell types as well.

To further investigate secretory cell type specific signatures of the enteroids, we measured enrichment of secretory lineages in the upregulated DEG lists using additional marker genes of Paneth cells, goblet cells and enteroendocrine cells. While all tests were significant (hypergeometric model, q value ≤ 0.05), we identified greater enrichment of Paneth cell markers in the PCeE DEG list and goblet markers in the GCeE DEG list (Figure 2E). This confirms that both enteroid enrichment protocols were successful in increasing the proportion of their target cell type, but also increased proportions of other secretory lineages, albeit to a lesser extent (Table S4). This observation confirms previous studies that these enteroid differentiation protocols result in enteroendocrine enrichment in addition to Paneth cell and goblet cell enrichment (25,28).

In conclusion, we have used image-based validation, pathway analysis and marker gene investigation to show successful enrichment of target cell types in the PCeE’s and GCeE’s. We also highlighted an additional increase in other secretory lineages, particularly enteroendocrine cells, as a result of both enrichment protocols.

### Reconstruction of regulatory networks altered by enteroid differentiation skewing

To gain an understanding of the regulatory changes occuring when enteroids development is skewed, we applied a network biology approach, and identified regulator-target relationships within the DEG lists. First, we generated a large network of non-specific molecular interactions known to occur in mice, by collating lists of published data (Table S6). The resulting network (termed the universal network) consisted of 1,383,897 unique regulatory interactions connecting 23,801 molecular entities. All interactions within the network represent one of the following types of regulation, where every node is a DEG: TF-TG, TF-lncRNA, TF-miRNA, miRNA-mRNA or lncRNA-miRNA. TF-TGs and TF-lncRNAs make up the majority of the network at 77% and 11% of all interactions, respectively. Due to its non-specific nature, this universal network contains many interactions not relevant for the current biological context. In order to get a clearer and valid picture of regulatory interactions occurring in our enteroids, we used the universal network to annotate the PCeE DEGs and GCeE DEGs with regulatory connections. Combining these connections, we generated specific regulatory networks for PCeEs and GCeEs, where every node is a DEG and every interaction has been observed in mice previously.

In total, the PCeE network, generated using differential expression data from the PCeEs compared to the conventionally differentiated enteroids, contained 37,062 interactions connecting 208 unique regulators with 3,023 unique targets (Figure 3A, Item S1). The GCeE network, generated using differential expression data from the GCeEs compared to the conventionally differentiated enteroids, contained 19,171 interactions connecting 124 unique regulators with 2,095 unique targets (Figure 3A, Item S1). 15.7% of all interactions (8,856 out of 56,234) were shared between the PCeE and GCeE networks, however the interacting molecular entities in these interactions (termed nodes) did not all exhibit the same direction of differential expression between the networks (comparing PCeE or the GCeE data to the conventionally differentiated enteroid data). In each of the enriched enteroid regulatory networks, a particular gene was represented (as a node in the network) only once, but may have been involved in multiple different interactions. In different interactions, a single node could act either as a regulator or as a target and in different molecular forms, for example, as a lncRNA in one interaction and as a target gene in another.

**Figure 3:**
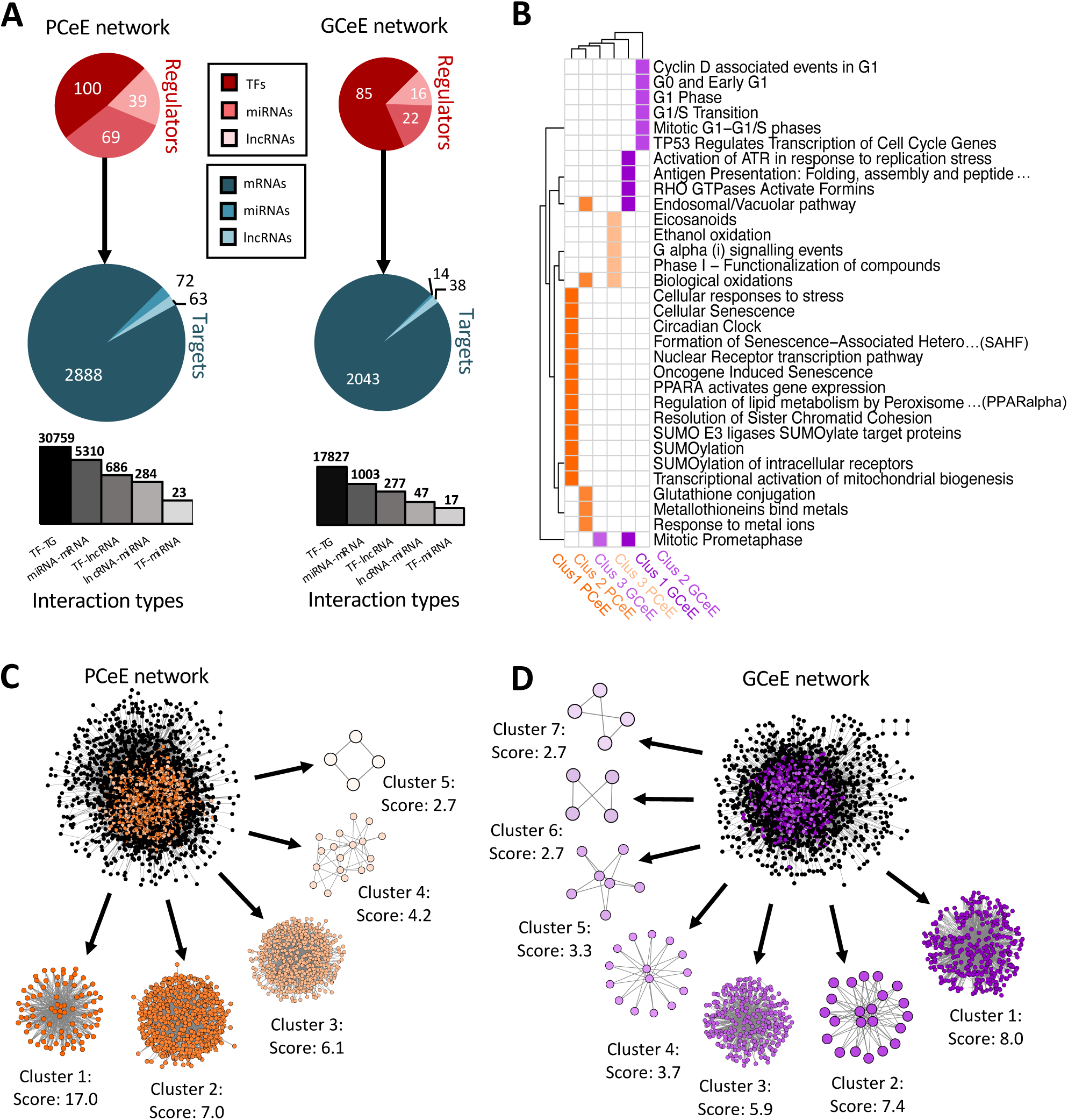
Summary and cluster analysis of regulatory network for Paneth cell enriched enteroid (PCeE) and goblet cell enriched enteroid (GCeE) datasets. **A:** Summary of number of nodes and interactions in the whole PCeE (left) and GCeE (right) networks. Total number of each regulator type shown in red, number of each target type shown in blue. In the targets pie-chart, mRNAs also represent protein coding genes and proteins, miRNAs also represent miRNAs genes and lncRNAs also represent lncRNA genes. Size of circles represents log10(total unique regulators/targets). Bar chart represents the distribution of interaction types in the networks (log10 scale). **B:** Heatplot of Reactome pathways significantly associated (q val ≤ 0.05) with each cluster of the PCeE (orange) and GCeE (purple) networks. Only the top 5 pathways shown for each group (or more where equal q values). Only the top 3 clusters had significantly associated pathways. Clusters labelled with rank and cell type and colours match the colour of the cluster shown in C and D. **C, D :** Visualisation of the PCeE and GCeE regulatory networks with their associated clusters. The cluster rank and score is given next to each cluster. Black nodes in the whole networks represent nodes which were not found in any cluster, whereas coloured nodes represent the cluster which they are part of. TF-transcription factor; miRNA-microRNA; lncRNA-long non-coding RNA. See Items S1, S2 and Table S5.

To further investigate the makeup of these networks, we employed cluster analysis to identify highly interconnected regions (possible regulatory modules) in the PCeE and GCeE regulatory networks. Using the MCODE software (37), we identified five distinct clusters in the PCeE network and seven distinct clusters in the GCeE networks. A total or 1314 nodes are present in the PCeE network clusters and 698 in the GCeE network clusters. Functional analysis identified Reactome pathways (38) associated with each of the modules. Significant pathways (q val ≤ 0.05) were identified only for the highest ranked three modules from each network, with a total of 12 pathways shared between the PCeE and GCeE associated clusters (out of 32 associated with the PCeE clusters and 42 with the GCeE clusters) (Figure 3B-D). Of particular note, the first cluster of the GCeE network has associations with the endosomal/vacuolar pathway and antigen presentation, the second cluster is associated with the cell cycle. Of the PCeE clusters, the first cluster is associated with a range of functions including nuclear receptor transcription pathway, regulation of lipid metabolism and senescence. The second is associated with response to metal ions and endosomal/vacuolar pathway and the third with G alpha (i) signalling events (Table S5).

In conclusion, we have generated regulatory interaction networks, including transcriptional and post-transcriptional interactions, which illustrate the effect of skewing enteroid differentiation towards Paneth cell and goblet cell lineages.

### Identification of potential cell type specific master regulators

Through pathway and marker analysis we predicted that our PCeE and GCeE datasets (i.e. DEG lists), and consequently our regulatory networks, contain signatures from the cell type of interest as well as additional noise from other secretory lineages. To focus specifically on the cell type specific elements of the networks, we used previously identified cell type specific markers to extract predicted Paneth cell and goblet cell regulators from our PCeE and GCeE networks. As cell type specific markers represent genes performing functions specific to a particular cell type, we expected that the regulators of these marker genes will have an important role in determining the function of said cell type. To identify these regulators, we extracted from the PCeE and GCeE networks, all relevant cell type specific markers and their direct regulators. These new networks were termed the Paneth cell subnetwork and goblet cell subnetwork respectively. The Paneth cell subnetwork contained 33 markers specific for Paneth cells with 62 possible regulators. The goblet cell subnework contained 150 markers with 63 possible regulators (Figure 4, Table S7). Observing the ratio of regulators and markers, the Paneth cell subnetwork had, on average, 1.88 regulators for each marker. On the other hand, the goblet cell subnetwork exhibited only 0.42 regulators for each marker. The quantity of markers identified in each subnetwork (33 in the Paneth network and 150 in the goblet network) correlates with the number of marker genes identified by Haber *et al.* (12). However, far fewer regulators were identified in the goblet cell subnetwork per marker than for the Paneth cell subnetwork. Whilst the underlying reason for this discrepancy is unknown, it could potentially be evidence of the complex regulatory environment required to integrate and respond to the arsenal of signals recognised by Paneth cells in comparison to goblet cells (35).

**Figure 4:**
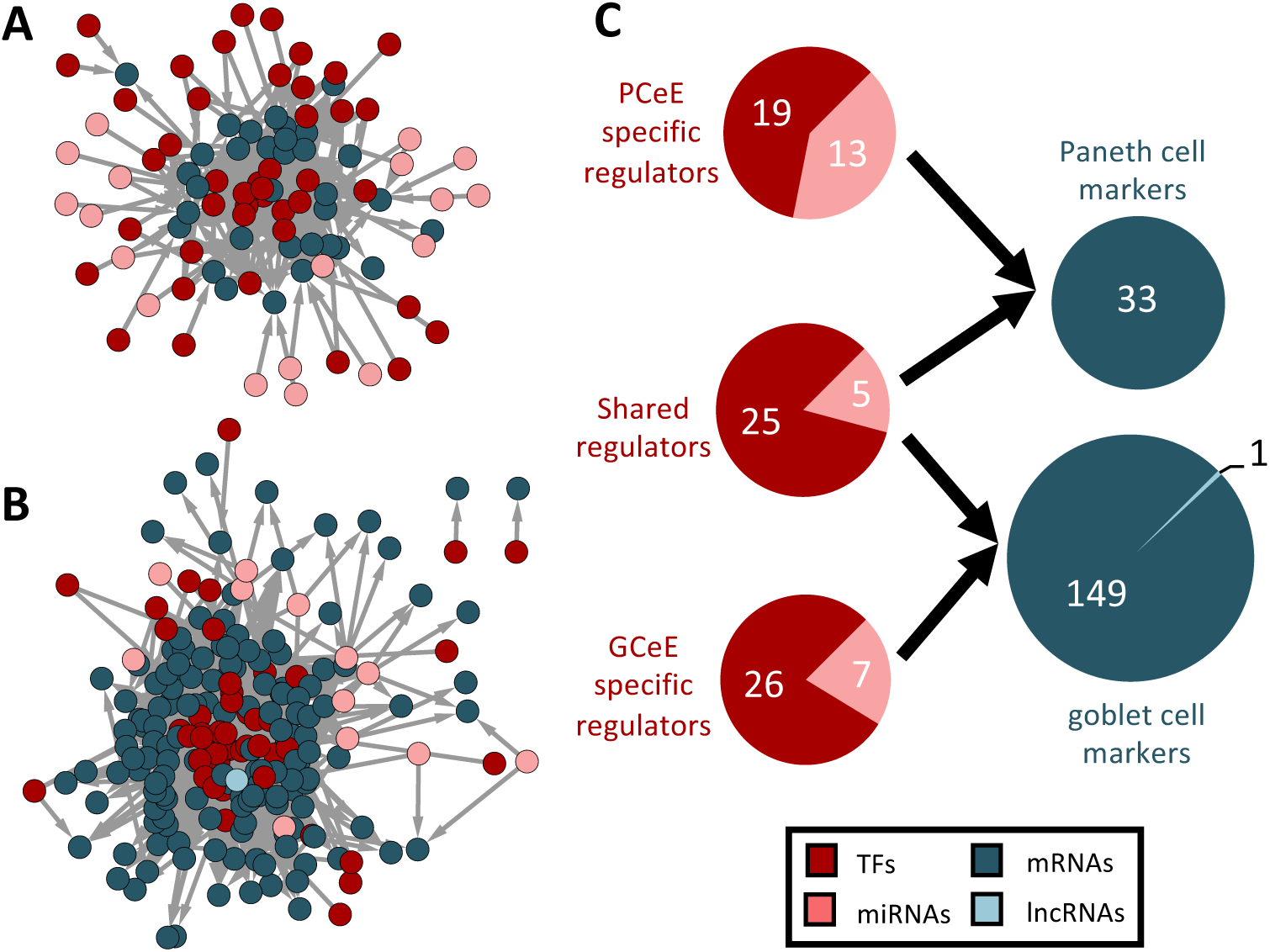
Regulator-marker subnetworks for Paneth cell and goblet cell datasets. **A, B:** Paneth cell (A) and goblet cell (B) subnetworks. Nodes represent genes, transcription factors or RNAs and edges represent directed physical regulatory connections. Regulators are shown in red and pink. Cell type specific markers are shown in blue. **C:** Summary of the number of nodes present in both the subnetworks. Paneth cell data above and goblet cell data below. Total number of each regulator type shown in red, number of each target type shown in blue. Regulators have been categorised based on their membership in the two subnetworks - shared regulators are present in both networks. In the targets pie-chart, mRNAs also represent protein coding genes. Size of circles represents log10 (total unique regulators/targets). TF-transcription factor; miRNA- microRNA; lncRNA-long non-coding RNA.

Of the 95 marker regulators, we identified approximately one-third (30/95) as present in both subnetworks (Figure 4C). Given that the markers are different between the cell types, a regulator shared between the Paneth cell and goblet cell subnetworks must show an altered pattern of regulatory targeting in the two cell types. This phenomenon, referred to as regulatory rewiring, often results in functional differences of shared regulators in different environments (39) - for example, in this case, between the Paneth cells and goblet cells.

Further investigation of the distinct regulator-marker interactions highlighted a gradient of regulator specificity. We generated matrices to visualise the markers controlled by each regulator in the goblet cell (Figure 5A) and the Paneth cell (Figure 5B,C) subnetworks. Each coloured square indicates that a marker (shown on the y-axis) is regulated by the corresponding regulator (shown on the x-axis). Squares are coloured green if the associated regulator is shared between the Paneth cell and goblet cell subnetworks and orange if they are specific to one subnetwork. A collection of regulators (both subnetwork specific and shared) appear to regulate large proportions of the markers. For example, Ets1, Nr3c1 and Vdr regulate >50% of the markers in both the Paneth cell and the goblet cell subnetworks. Specific to the Paneth cell subnetwork, Cebpa, Jun, Nr1d1 and Rxra regulate >50% of the markers. Specific to the goblet cell subnetwork, Gfi1b and Myc regulate >50% of the markers. These regulators represent potential master regulators of differentiation or maintenance of the given cell types in the enriched enteroids. Referring back to the highly-interconnected clusters identified in the PCeE and GCeE networks (Figure 3C-D), we find these predicted master regulators in different clusters. In the PCeE network, Cebpa, Nr1d1, Nr3c1 and Rxra are in cluster 1, Vdr is in cluster 2, Jun is in cluster 3 and Ets1 is unclustered. In the GCeE network, Ets1 and Myc are in cluster 1, Nr3c1 and Vdr are in cluster 2 and Gfi1b is in cluster 3. This suggests a wide range of central functions are carried out by this group of regulators, with possible divergence of roles between the Paneth cell and the goblet cell. In contrast to the predicted master regulators, regulators such as Mafk in the Paneth cell subnetwork and Spdef in the goblet cell subnetwork regulate only one marker. These regulators likely have more functionally specific roles.

**Figure 5:**
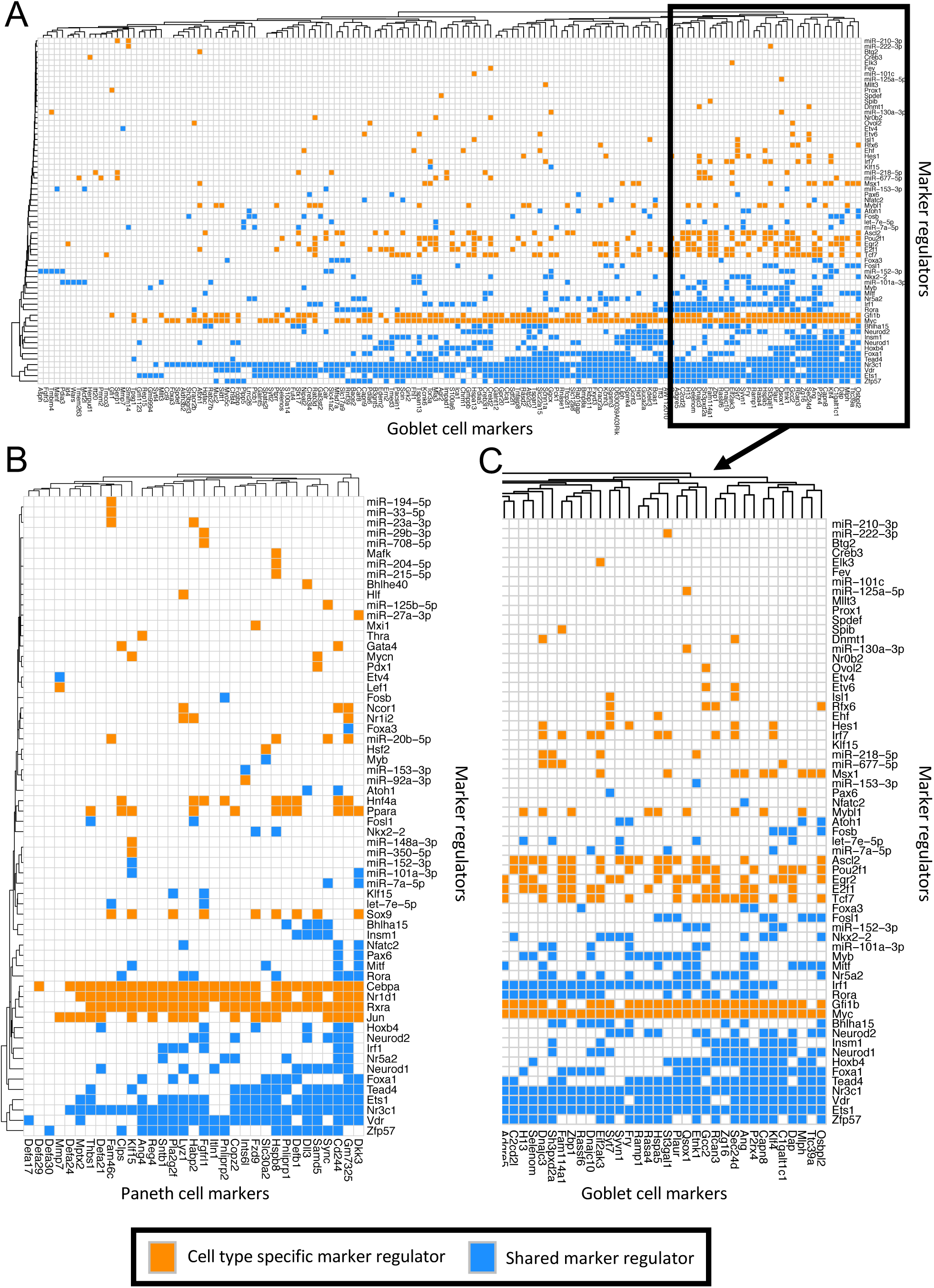
Matrices of interactions between markers and their regulators in the Paneth cell and goblet cell subnetworks. Regulators on y-axis, markers (regulator targets) on y-axis. Orange boxes indicate the interaction of a regulator and a marker where the regulator is only found in one of the two subnetworks. Blue boxes signify that the regulator is found in both the Paneth cell and the goblet cell subnetworks. **A:** All goblet cell markers (12) and their regulators in the goblet cell subnetwork. **B:** All Paneth cell markers (12) and their regulators in the Paneth cell subnetwork. **C:** Sub-section of A showing the markers (and their regulators) which have the most regulatory connections. Interactions in Table S7.

Together, these results highlight potential regulators which likely play key roles in specification and maintenance of Paneth and goblet cells and their functions in cell type enriched enteroids.

### Regulators of cell type specific markers exhibit rewiring between Paneth cells and goblet cells

Cell type specific markers, which carry out cell type specific functions, are inherently different between the Paneth cell and goblet cell subnetworks (mutually exclusive). Therefore, the regulators observed in both Paneth cell and goblet cell subnetworks (shared regulators) are expected to target different marker genes. To do this, the regulators must have different regulatory connections in the different cell types, a phenomenon termed ‘rewiring’ (40). We extended the analysis to the original regulatory networks (PCeE and GCeE networks) to investigate whether any of the 30 identified shared regulators are rewired between the whole PCeE and the GCeE networks, and thus are highly likely to have different functions in the two types of enriched enteroids as well as between Paneth and goblet cells. To quantify rewiring of each of these regulators, we observed their targets in the PCeE and GCeE networks using the Cytoscape application, DyNet. DyNet assigns each regulator a rewiring score depending on how different their targets are between the two regulatory networks (Table S8). Using these rewiring scores, we identified the five most rewired regulators (of 30) as Etv4, let-7e-5p, miR-151-3p, Myb and Rora. Functional enrichment analysis was carried out on the targets of these regulators to test whether the targets specific to the PCeE and GCeE networks have different functions (hypergeometric model, q value ≤ 0.1) (Table S9). Across all five regulators the general trend indicated that targets specific to the PCeE network are associated with metabolism; targets specific to the GCeE network are associated with cell cycle and DNA repair. As pathway analysis carried out on the enteroid DEGs (Figure 2D) identified the same phenomenon, this suggests that the rewired regulators could be key drivers of transcriptional changes during the skewing of enteroid differentiation towards Paneth cell or goblet cell lineages. In addition, given that the strongest signal of enriched enteroids represents their enriched cell type, we predict that these functions are key features of Paneth cells and goblet cells in the enteroids, and that the rewired regulators are important drivers of cell type specific functions.

Looking at the regulators in more detail, the GCeE specific targets of miR-151-3p, for example, are significantly enriched in functions relating to antigen presentation, cell junction organisation, Notch signalling and the calnexin/calreticulin cycle. None of these functions are enriched in the shared or PCeE targets. Of particular interest is the calnexin/calreticulin cycle, which is known to play an important role in ensuring proteins in the endoplasmic reticulum are correctly folded and assembled (41). Dysfunction of protein folding and the presence of endoplasmic reticulum stress are both associated with IBD (42–44). Therefore, we predict that miR-151-3p plays a role in the secretory pathway of goblet cells and could be an interesting target for IBD research. In addition, different functional profiles were also observed for the targets of Rora in the PCeE and GCeE regulatory networks: targets present in both networks are significantly associated with mitosis, whereas those specific to the PCeE network are associated with metabolism, protein localisation, nuclear receptor transcription pathway, circadian clock and hypoxia induced signalling. GCeE specific targets of Rora are connected to Notch signalling, cell cycle and signalling by Rho GTPases (associated with cell migration, adhesion and membrane trafficking) and interferon.

Altogether these observations show that some of the regulators of both Paneth cell and goblet cell marker genes have different targets (with different associated functions) between the PCeE and the GCeE networks. This suggests that regulatory rewiring occurs between Paneth cell and goblet cell types.

### Evaluating the disease relevance of the subnetwork specific master regulators

To investigate the function and relevance of the predicted master regulators in IBD, we carried out three analyses: 1) a literature search to check what is known about the identified master regulators; 2) an enrichment analysis to evaluate the disease relevant genes in the PCeE and GCeE networks and among the targets of the predicted master regulators; and finally, 3) a comparative analysis with human biopsy based single cell dataset to confirm the relevance of the data we identified PCeE and GCeE networks.

The literature search was carried out using the three groups of predicted master regulators: those specific to the Paneth cell markers (Cebpa, Jun, Nr1d1 and Rxra), those specific to the goblet cell markers (Gfi1b and Myc) and those which appear to regulate many of the markers of both cell types (Ets1, Nr3c1 and Vdr). We identified five genes (*Ets1, Nr1d1, Rxra, Nr3c1 and Vdr*) with associations to inflammation, autophagy and/or inflammatory bowel disease (IBD), as shown in Table 1. These genes correspond to 71% (5/7) of the Paneth cell associated master regulators and 60% (3/5) of the goblet cell associated master regulators. Interestingly, four of these genes (all apart from *Ets1*), encode nuclear hormone receptors.

**Table 1:** Literature associations relating to autophagy, inflammation and IBD for putative master regulators.

| Putative master regulator | Autophagy / inflammation / IBD associations | References |
| --- | --- | --- |
| NR1D1 (REV-ERBa) | Modulates autophagy and lysosome biogenesis in macrophages leading to antimycobacterial effects | (45) |
|  | SNP rs12946510 which has associations to IBD, acts as a cis-eQTL for NR1D1 | (46) |
| NR3C1 (glucocorticoid receptor) | Associations with cellular proliferation and anti-inflammatory responses | (47) |
|  | Exogenous glucocorticoids are heavily used as anti-inflammatory therapy for IBD | (48,49)) |
|  | ATG16L1, an autophagy related gene, was down-regulated in patients who do not respond to glucocorticoid treatment | (50,51) |
|  | Transcriptionally regulates NFKB1, a SNP affected gene in ulcerative colitis | (52,53) |
| VDR (Vitamin D Receptor) | Regulates autophagy in Paneth cells through ATG16L1 – dysfunction of autophagy in Paneth cells has been linked to Crohn's disease | (54,55) |
|  | Induces antimicrobial gene expression in other cell lines | (56,57) |
|  | Specific polymorphisms in the VDR genes have been connected to increased susceptibility to IBD | (58) |
|  | A study looking at colonic biopsies of IBD patients observed reduced VDR expression compared to healthy biopsies | (59) |
|  | Interacts with five SNP affected UC genes | (60,61) |
| RXRα (Retinoid X Receptor Alpha) | Heterodimerizes with VDR (see above) | (62) |
| ETS1 (ETS Proto-Oncogene 1) | Important role in the development of hematopoietic cells and Th1 inflammatory responses | (63,64) |
|  | Angiogenic factors in the VEGF-Ets-1 cascades are upregulated in UC and downregulated in CD | (65) |
|  | IBD susceptibility gene | (66) |

Given the possible relationship between the identified master regulators and IBD, we tested the potential of the PCeE and GCeE regulatory networks to study the pathomechanisms of CD or UC. We checked for the presence of known CD or UC associated genes in the networks, using data from two studies of single nucleotide polymorphisms (SNPs) (67,68) and one study of CD expression quantitative trait loci (eQTLs) (69). Using hypergeometric significance tests, we found that the PCeE network was significantly enriched in all tested lists: Genes with UC associated SNPs (13/47, p < 0.005), genes with CD associated SNPs (22/97, p < 0.005) and genes with CD associated eQTLs (290/1607, p < 0.0001) (Table S10-12). On the other hand, we found that the GCeE network was significantly enriched in genes with UC associated SNPs (10/47, p < 0.005) but regarding CD, the genes with SNP associations were not significantly enriched (12/97, p = 0.11) and the genes with qQTL associations were enriched with a larger p value (p < 0.05) (Table S10-12).

Next we investigated whether any of the genes with UC or CD associated SNPs acts as regulators in the PCeE and GCeE networks. Of the genes with CD associated SNPs, one acts as a regulator in each network. Similarly, two genes with UC associated SNPs act as regulators in the networks. Specifically regarding CD associated genes, in the PCeE network, the gene *Dbp* regulates *Bik*, which encodes the BCL2 interacting killer, a pro-apoptotic, death promoting protein. In the GCeE network, *Notch2* regulates *Notch3* and *Hes1.* Specifically, regarding UC associated genes, in the PCeE network, *Hnf4a* regulates 994 genes/RNAs including nine Paneth cell markers (*Cd244a, Fgfrl1, Clps, Habp2, Hspb8, Pnliprp1/2, Defb1, Mymx*) and one other gene with UC associated SNPs (*Tnfsf15*). Additionally, a gene with UC associated SNPs, *Nr5a2*, was found in both the PCeE and GCeE networks regulating 389 and 276 genes/RNAs respectively. In the PCeE network *Nr5a2* targets include 6 Paneth cell markers (*Cd244a, Copz2, Pnliprp1/2, Sntb1, Mymx*). Ultimately, the large number of targets of these regulatory UC associated genes suggests they have wide ranging effects on the regulatory network of Paneth and goblet cells. To further establish the relevance of the inferred PCeE and GCeE networks, we also found an over-representation of drug target associated genes in both the PCeE and GCeE networks (2683/16223 and 1918/16223 respectively, p < 0.0001), highlighting their potential for the study of therapeutic implications (Table S12).

To investigate the link between predicted master regulators and IBD, we observed whether the genes with UC and CD associated SNPs are regulated by the predicted master regulators in the PCeE and GCeE networks (Table S13). Given that Paneth cell dysregulation is classically associated with CD and goblet cell dysregulation/depletion with UC (70), we focused this analysis only on these pairings, examining CD genes amongst targets of Paneth cell predicted master regulators, and UC genes amongst targets of goblet cell predicted master regulators. In the PCeE network, we found 21 (of 22) of the CD genes were regulated by at least one of the seven Paneth cell predicted master regulators, while the targets of these master regulators were significantly enriched with CD genes in the PCeE network (p < 0.001). Similarly, we observed that all 10 UC genes in the GCeE network were regulated by at least one of the five goblet cell predicted master regulators, while the targets of these master regulators were significantly enriched with UC genes (p < 0.005).

To confirm the relevance of these predicted master regulators in a human system, a similar analysis was carried out using goblet cell differentially expressed genes from a recent single cell study of human inflamed UC colon biopsies (13). Using the top 100 differentially expressed genes, following conversion to mouse Ensembl identifiers, 20 were found to be targeted by the predicted goblet cell master regulators in the GCeE network. This represents a significant enrichment amongst all master regulator targets (p < 0.005) (Table S13).

Ultimately, by integrating functional annotations obtained through literature searches, we show that the Paneth cell and goblet cell regulatory networks contain genes with direct and indirect associations with IBD. Furthermore, we find that the PCEe and GCeE networks and the targets of predicted master regulators are enriched with IBD associated genes - this finding is corroborated using human single cell data from UC colon biopsies. Consequently, these networks and the workflow to reconstruct and analyse them have great potential for the study of IBD pathomechanisms in specific intestinal cell types.

## Discussion

By generating and integrating cell type enriched enteroid RNAseq datasets with regulatory networks, we characterised the regulatory environment altered by differentiation skewing. Through focusing on cell type specific markers, we used the regulatory networks to predict master regulators of Paneth cells and goblet cells and to highlight the role of regulatory rewiring in cell differentiation. Given the relevance of Paneth cell and goblet cell dysfunction in IBD, future application of cell type enriched enteroids combined with our network analysis workflow can be used to disentangle multifactorial mechanisms of IBD.

Analysis using known cell type specific markers confirmed that skewing differentiation of enteroids towards Paneth cells or goblet cells results in an increase in the target cell signatures at the transcriptomics level. In addition, signatures of enteroendocrine cells were increased in both differentiation protocols, albeit at a lower amount than the target cell type. This finding correlates with previous investigations at both the transcriptomic and proteomic levels (25,27–29) and is likely due to the shared differentiation pathways of these secretory cells. Nevertheless, as enteroids contain a mixed population of cell types by nature and because intercellular communication is key to a functioning epithelium (19,71), the increased proportion of non-targeted secretory lineages should not be an issue for the application of these models to research. In fact, the enrichment of specific cell types is beneficial for enteroid-based research to increase the signal originating from a specific population of cells and to provide a larger population of cells of interest for downstream single cell analysis of enteroids, which is particularly beneficial when studying rare populations such as Paneth cells. This is valuable due to the lack of *in vitro* models for long-term culture of non-self-renewing small intestinal epithelial cells (72,73). Specifically, the comparison of ‘omics data from a cell type enriched enteroid to a conventionally differentiated enteroid enables generation of cell type signatures with more specificity than can be obtained otherwise - except through single cell sequencing. Single cell sequencing, however, comes at a greater financial cost and provides lower coverage which can be problematic for rare cell types and lowly expressed RNAs. Furthermore, a number of previous studies have shown that these cell type enriched enteroid models, which offer a simplified and manipulatable version of the intestinal environment, are useful for the investigation of health and disease related processes (26,28,29). It must be considered, however, that through applying chemical inhibitors to enteroids to enrich particular cell types, we may observe changes in gene expression which are related to the direct effects of the inhibitor but not to differentiation. Due to the nature of the inhibitors, it would be challenging to separate these effects. For example, CHIR99021, which is used for enriching Paneth cells, is a direct inhibitor of glycogen synthase kinase 3 (GSK-3). GSK-3 has a role in many cellular pathways including cellular proliferation and glucose regulation (74).

Using *a priori* molecular interaction knowledge, we annotated differentially expressed genes (cell type enriched enteroids vs. conventionally differentiated enteroids) with regulatory connections. Collecting all connections together generated regulatory networks for the Paneth cell enriched enteroid (PCeE) and goblet cell enriched enteroid (GCeE) datasets. This approach to collating networks (regulatory or otherwise) has been used for a wide variety of research aims, such as the identification of genes functioning in a variety of diseases (75,76), the prioritisation of therapeutic targets (77) and for a more general understanding of gene regulation in biological systems (78,79). The application of prior knowledge avoids the need for reverse engineering / inference of regulatory network connections, which is time consuming, computationally expensive and requires large quantities of high quality data (80). To investigate the substructure and functional associations of the generated PCeE and GCeE regulatory networks we applied a clustering approach. The identified clusters represent collections of highly interconnected nodes, which likely form regulatory modules. Functional analysis confirmed distinct functional associations between the clusters as well as between the networks. The observation that less than half of the network nodes exist in clusters is consistent with the view that regulatory networks are hierarchical and scale free with most genes exhibiting low pleiotropy (81,82).

Given the observed additional increase in secretory lineages based on the DEGs of the enriched organoids, we chose to use cell type specific markers to extract interactions specific to Paneth cells and goblet cells from the generated regulatory networks. This enables further enrichment of Paneth cell and goblet cell signatures and reduction in noise in the networks due to the presence of other cell types in the enteroids. Using this approach, we identified possible regulators of cell type specific functions in Paneth cells and goblet cells. Some of these regulators were predicted to be important in both cell types, but exhibited differential targeting patterns between the PCeE and the GCeE networks, indicating rewiring of regulators between the cell types. This highlights apparent redundancy and/or cooperation of regulators which control similar cell type specific functions and shows the potential importance of regulatory rewiring in the evolution of cell type specific pathways and functions, something which has been shown previously to occur (83,84). Functional analysis of the targets of the most rewired regulators (Etv4, let-7e-5p, miR-151-3p, Myb and Rora) highlights an overrepresentation of metabolism associated targets in the PCeE network and cell cycle associated targets in the GCeE network. A similar result was observed when functional analysis was carried out on genes with significantly different expression levels between the cell type enriched enteroids and the conventionally differentiated enteroids (Figure 2B). This suggests that transcriptional changes during the skewing of enteroid differentiation could be driven by rewired regulators and that these functions are key features of Paneth cells and goblet cells in the enteroids. The latter is supported by current understanding that Paneth cells rely on high levels of protein and lipid biosynthesis for secretory functions (85)important role in metabolically supporting stem cells (86). Additionally, as terminally-differentiated cells do not undergo cell division, this result suggests that enteroid goblet cell signatures are derived from a large population of semi-differentiated goblet-like cells, a phenomenon previously observed in tissue sample based studies (13,87). Extension of our workflow to single cell sequencing of enteroid cells could validate these findings by providing greater cell type specificity. However, these techniques pose further technical and economic challenges (88,89). Specifically, a large number of organoids must be sequenced to mitigate cellular complexity and batch heterogeneity and powerful, reproducible and accurate computational pipelines are required to analyse such data (90).

We predicted key regulators involved in differentiation or maintenance of Paneth cells and goblet cells in the enteroids: Cebpa, Jun, Nr1d1 and Rxra specific to Paneth cells, Gfi1b and Myc specific for goblet cells and Ets1, Nr3c1 and Vdr shared between them. Validation of these regulators poses significant challenges due to their wide expression and broad function range. If the master regulators are controlling differentiation as opposed to cell function maintenance, evaluating lineage arrest or delay can be carried out using a gene knockout or knock down. However the effects of pleiotropy will significantly hamper the results and such a study would require significant follow-up studies. On the other hand, if key regulators were predicted by applying the presented computational workflow to condition-specific organoids compared to control organoids (eg. drug treated organoids vs non-treated organoids), the validation would be much simpler. Literature investigation highlighted that many of the predicted master regulators, particularly those associated with Paneth cells, have connections to autophagy, inflammation and IBD (Table 1). While some of these associations are related to different cell types, we assume that if a gene can contribute to a specific function in one cell type it may also contribute in another. This finding suggests that dysregulation of key cell master regulators could contribute to IBD.

To further investigate this finding, we identified Crohn’s disease (CD) and ulcerative colitis (UC) genes in the PCeE and GCeE networks. We found that CD associated genes are more strongly associated with the PCeE network than the GCeE network. Given that Paneth cell dysfunction is classically associated with CD, this finding highlights the relevance of the generated networks to the *in vivo* situation. In the PCeE network one SNP associated CD gene, *Dbp*, acts as a regulator. *Dbp*, encoding the D site binding protein, regulates *Bik*, which encodes the BCL2 interacting killer, a pro-apoptotic, death promoting protein. Interestingly, rate of apoptosis has been implicated in IBD disease mechanisms (91) and has been associated with IBD drug response (92). Therefore, this finding highlights a possible regulatory connection between CD susceptibility genes and IBD pathology on a Paneth cell specific level. In the GCeE network, the SNP associated CD gene *Notch2* acts as a regulator for *Notch3* and *Hes1.* It has been previously demonstrated that this pathway can block glucocorticoid resistance in T-cell acute lymphoblastic leukaemia via NR3C1 (predicted master regulator) (93). This is relevant to IBD given that glucocorticoids are a common treatment for IBD patients (49). Furthermore, this pathway has been previously associated with goblet cell depletion in humans (94) commonly observed in IBD patients. Furthermore, we identified a significant enrichment of UC associated genes in both the PCeE and GCeE networks. The majority of UC associated genes identified in the networks (9/14) were present in both, suggesting that genetic susceptibilities of UC do not have a Paneth cell or goblet cell specific effect. Two of the identified UC associated genes act as regulators in the networks (*Nr5a2* and *Hnf4a)*, targeting hundreds of genes and thus suggesting a broad ranging effect on the networks. Building on the identified literature associations of predicted master regulators, we found that the targets of Paneth cell master regulators are enriched with CD associated genes, and the targets of the goblet cell master regulators are enriched with UC associated genes. This finding was further illustrated using UC associated goblet cell genes from a human biopsy study (13), highlighting the relevance of these findings in a human system. Ultimately, the observation of IBD susceptibility genes in the regulatory networks of these enteroids highlights possible application of this model system to study disease regulation in specific intestinal cell types, through understanding specific mechanistic pathways.

We have shown how network biology techniques can be applied to generate interaction networks representing the change in regulatory environments between two sets of enteroids. Here we presented this workflow, and by using transcriptomics data we characterised the effect of Paneth cell and goblet cell differentiation skewing protocols on enteroids. However. the described workflow could be applied to a variety of ‘omics datasets and enteroid conditions. For example, to test the response of enteroids to external stimuli, such as bacteria, and on enteroids grown from human-derived biopsies, enabling patient-specific experiments. The application of further ‘omics data-types to the described approach could generate a more holistic view of cellular molecular mechanisms, including the ability to observe post-translational regulation. In this study, we integrated miRNA and lncRNA expression datasets, in addition to mRNA data, as these molecules have been shown to perform critical regulatory and mediatory functions in maintaining intestinal homeostasis (46,68,95). However, only small proportions of the generated PCeE and the GCeE networks contained miRNA and lncRNA interactions (Figure 2B), due to lack of published interaction information, particularly from murine studies. Both the application of human enteroid data and the future publication of high-throughput interaction studies involving miRNAs and lncRNAs will improve the ability to study such interactions. Nevertheless, these network approaches are beneficial for contextualising gene lists through annotating relevant signalling and regulatory pathways (31) and we can use them to represent and analyse current biological knowledge, to generate hypotheses and to guide further research.

In conclusion, we described an integrative systems biology workflow to compare regulatory landscapes between enteroids from different conditions, incorporating information on transcriptional and post-transcriptional regulation. We applied the workflow to compare Paneth cell and goblet cell enriched enteroids to conventionally differentiated enteroids and predicted Paneth cell and goblet cell specific regulators, which could provide potential targets for further study of IBD mechanisms. Application of this workflow to patient derived organoids, genetic knockout and/or microbially challenged enteroids, alongside appropriate validation and single cell sequencing if available, will aid discovery of key regulators and signalling pathways of healthy and disease associated intestinal cell types.

## Supporting information

Tabel S12

## Acknowledgement

The authors are grateful for the helpful discussions to the past and present members and visitors of the Wileman, Haerty and Korcsmaros groups.

## Author contributions

EJJ, ZJM and IH worked on the transcriptomics pipeline development; AT, PS, TWr, JB and MO designed and carried out specific network analysis tasks; SRC, UM, PPP and TWi established and supervised the organoid enrichment work; LJH and FDP supervised the experimental and transcriptomics analysis work; WH, TWi and TK designed the study, supervised the experiments and the analysis. AT, PS, ZM, EJ, IH and TK wrote the manuscript.

## Methods

### Animal handling

C57BL/6J mice of both sexes were used for enteroid generation. All animals were maintained in accordance with the Animals (Scientific Procedures) Act 1986 (ASPA).

### Small intestinal organoid culture

Murine enteroids were generated as described previously (18,20,29). Briefly, the entire small intestine was opened longitudinally, washed in cold PBS then cut into ∼5mm pieces. The intestinal fragments were incubated in 30mM EDTA/PBS for 5 minutes, transferred to PBS for shaking, then returned to EDTA for 5 minutes. This process was repeated until five fractions had been generated. The PBS supernatant fractions were inspected for released crypts. The crypt suspensions were passed through a 70μm filter to remove any villus fragments, then centrifuged at 300xg for 5 minutes. Pellets were resuspended in 200μl phenol-red free Matrigel (Corning), seeded in small domes in 24-well plates and incubated at 37°C for 20 minutes to allow Matrigel to polymerise. Enteroid media containing EGF, Noggin and R-spondin (ENR; (18)) was then overlaid. Enteroids were generated from three separate animals for each condition, generating three biological replicates.

To chemically induce differentiation, on days two, five and seven post-crypt isolation, ENR media was changed to include additional factors for each cell type specific condition: 3μM CHIR99021 (Tocris) and 10μM DAPT (Tocris) [Paneth cells]; 2μM IWP-2 (Tocris) and 10μM DAPT [goblet and enteroendocrine cells] (25).

### Small intestinal organoid immunofluorescence

On day eight post-crypt isolation, enteroids were fixed with 4% paraformaldehyde (PFA; Sigma-Aldrich) for 1 hour at 4°C prior to permeabilization with 0.1% triton X-100 (Sigma-Aldrich) and incubation in blocking buffer containing 10% goat serum (Sigma-Aldrich). Immunostaining was performed overnight at 4°C using primary antibodies: mouse anti-E-cadherin (BD Transduction Laboratories), rabbit anti-muc2 (Santa Cruz) and rabbit anti-lysozyme (Dako), followed by Alexa Fluor-488 and -594 conjugated secondary antibodies (ThermoFisher Scientific). DNA was stained with DAPI (Molecular Probes). Images were acquired using a fluorescence microscope (Axioimager.M2, equipped with a Plan-Apochromat 63x/1.4 oil immersion objective) and analysed using ImageJ/FIJI V1.51.

### RNA extraction

On day eight post-crypt isolation (allowing optimal cell type-enrichment as previously shown; Yin et al., 2014), enteroids were extracted from Matrigel (Corning, 356237) using Cell Recovery Solution (BD Bioscience, 354253), rinsed in PBS and RNA was extracted using miRCURY RNA Isolation Tissue Kit (Exiqon, 300115) according to the manufacturer’s protocol.

### Stranded RNA library preparation

The enteroid transcriptomics libraries were constructed using the NEXTflex™ Rapid Directional RNA-Seq Kit (Perkin Elmer, 5138-07) using the polyA pull down beads from Illumina TruSeq RNA v2 library construction kit (Illumina, RS-122-2001) with the NEXTflex™ DNA Barcodes – 48 (Perkin Elmer, 514104) diluted to 6µM. The library preparation involved an initial QC of the RNA using Qubit DNA (Life technologies, Q32854) and RNA (Life technologies, Q32852) assays as well as a quality check using the PerkinElmer GX with the RNA assay (CLS960010). Ligated products were subjected to a bead-based size selection using Beckman Coulter XP beads (A63880). As well as performing a size selection this process removed the majority of unligated adapters. Prior to hybridisation to the flow cell the samples were amplified to enrich for DNA fragments with adapter molecules on both ends and to amplify the amount of DNA in the library. The strand that was sequenced is the cDNA strand. The insert size of the libraries was verified by running an aliquot of the DNA library on a PerkinElmer GX using the High Sensitivity DNA chip (Perkin Elmer, CLS760672) and the concentration was determined by using a High Sensitivity Qubit assay and q-PCR. Libraries were then equimolar pooled and checked by qPCR to ensure the libraries had the necessary sequencing adapters ligated.

### Small RNA library preparation

The small RNA libraries were made using the TruSeq Small RNA Library Prep Kits (Illumina, 15004197), six-base indexes distinguish samples and allow multiplexed sequencing and analysis using 48 unique indexes ((Set A: indexes 1-12 (Illumina, RS-200-0012), Set B: indexes 13–24 (Illumina, RS-200-0024), Set C: indexes 25–36 (Illumina, RS-200-0036), Set D: indices 37–48 (Illumina, RS-200-0048)) (TruSeq Small RNA Library Prep Kit Reference Guide (Illumina, 15004197 Rev.G)). The TruSeq Small RNA Library Prep Kit protocol is optimised for an input of 1µg of total RNA in 5 µl nuclease-free water or previously isolated microRNA may be used as starting material (minimum of 10–50 ng of purified small RNA). Total RNA is quantified using the Qubit RNA HS Assay kit (ThermoFisher, Q32852) and quality of the RNA is established using the Bioanalyzer RNA Nano kit (Agilent Technologies, 5067-1511), it is recommended that RNA with an RNA Integrity Number (RIN) value ≥ 8 is used for these libraries as samples with degraded mRNA are also likely to contain degraded small RNA.

Library purification combines using BluePippin cassettes (Sage Science Pippin Prep 3% Cassettes Dye-Free (CDF3010), set to collection mode range 125-160bp) to extract the library molecules followed by a concentration step (Qiagen MinElute PCR Purification, 28004) to produce libraries ready for sequencing. Library concentration and size are established using HS DNA Qubit and HS DNA Bioanalyser. The resulting libraries were then equimolar pooled and qPCR was performed on the pool prior to clustering.

### Stranded RNA sequencing on HiSeq 100PE

The final pool was quantified using a KAPA Library Quant Kit (Roche, 07960140001), denatured in NaOH and combined with HT1 plus a 1% PhiX spike at a final running concentration of 10pM.The flow-cell was clustered using HiSeq PE Cluster Kit v3 (Illumina, PE-401-3001) utilising the Illumina PE HiSeq Cluster Kit V3 cBot recipe V8.0 method on the Illumina cBot. Following the clustering procedure, the flow-cell was loaded onto the Illumina HiSeq2000 instrument following the manufacturer’s instructions with a 101 cycle paired reads and a 7-cycle index read. The sequencing chemistry used was HiSeq SBS Kit v3 (Illumina, FC-401-3001) with HiSeq Control Software 2.2.68 and RTA 1.18.66.3. Reads in bcl format were demultiplexed based on the 6bp Illumina index by CASAVA 1.8, allowing for a one base-pair mismatch per library, and converted to FASTQ format by bcl2fastq.

### Small RNA sequencing on HiSeq Rapid 50SE

The final pool was quantified using a KAPA Library Quant Kit (Roche, 07960140001), denatured in NaOH and combined with HT1 plus a PhiX spike at a final running concentration of 20pM. The libraries were hybridized to the flow-cell using TruSeq Rapid Duo cBot Sample Loading Kit (Illumina, CT-403-2001), utilising the Illumina RR_TemplateHyb_FirstExt_VR method on the Illumina cBot. The flow-cell was loaded onto the Illumina HiSeq2500 instrument following the manufacturer’s instructions with a 51-cycle single read and a 7 cycle index read. The sequencing chemistry used was HiSeq SBS Rapid Kit v2 (Illumina, FC-402-4022) with a single read cluster kit (Illumina, GD-402-4002), HiSeq Control Software 2.2.68 and RTA 1.18.66.3. Reads in bcl format were demultiplexed based on the 6bp Illumina index by CASAVA 1.8, allowing for a one base-pair mismatch per library, and converted to FASTQ format by bcl2fastq.

### Differentially expressed transcripts

The quality of stranded reads was assessed by FastQC software (version 0.11.4) (96). All reads coming from technical repeats were concatenated together and aligned (in stranded mode, i.e. with ‘--rna-strandness RF’ flag) using HISAT aligner (version 2.0.5) (97). Subsequently, a reference-based de novo transcriptome assembly was carried out for each biological repeat and merged together using StringTie (version 1.3.2) with following parameters: minimum transcript length of 200 nucleotides, minimum FPKM of 0.1 and minimum TPM of 0.1 (98,99). Coding potential of each novel transcript was determined with CPC (version 0.9.2) and CPAT (version 1.2.2) (100,101). From the novel transcripts, only non-coding transcripts (as predicted by both tools) were included in final GTF file. Gene and transcript abundances were estimated with kallisto (version 0.43.0) (102). Sleuth (version 0.28.1) R library was used to perform differential gene expression (103). mRNAs and lncRNAs with an absolute log2 fold change of 1 and q value ≤ 0.05 were considered to be differentially expressed.

The small RNA reads were analysed using the sRNAbench tool within the sRNAtoolbox suite of tools (104). The barcodes from the 5’ end and adapter sequences from the 3’ end were removed respectively. Zero mismatches were allowed in detecting the adapter sequences with a minimum adapter length set at 10. Only reads with a minimum length of 16 and a read-count of 2 were considered for further analysis. The mice miRNA collection was downloaded from miRBase version 21 (105). The trimmed and length filtered reads were then mapped to the mature version of the miRBase miRNAs in addition to the annotated version of the mouse genome (version mm10). No mismatches were allowed for the mapping. A seed length of 20 was used for the alignment with a maximum number of multiple mappings set at 10. Read-counts normalised after multiple-mapping were calculated for all the libraries. The multiple-mapping normalised read-counts from the corresponding cell type enriched enteroids were compared against the conventionally differentiated enteroids to identify differentially expressed miRNAs in a pair-wise manner using edgeR (106). miRNAs with an absolute log2 fold change of 1 and false discovery rate ≤ 0.05 were considered to be differentially expressed.

Differentially expressed genes were grouped by their presence in the PCeE dataset, the GCeE dataset or in both. Each group of differentially expressed genes was tested for functional enrichment (hypergeometric model, q value ≤ 0.1) based on Reactome and KEGG annotations using the ReactomePA R package (38,107–109) following conversion from mouse to human identifiers using Inparanoid (v8) (110,111).

### Enrichment of marker genes

Cell type specific marker gene lists were obtained a mouse single cell sequencing survey (12). The cell type specific signature genes for Paneth, goblet and enteroendocrine cell types were obtained from the droplet-based and the plate-based methods. Gene symbols were converted to Ensembl gene IDs using bioDBnet db2db (112).

Hypergeometric distribution testing was carried out using a custom R script to measure enrichment of cell type specific marker genes in the differentially upregulated gene sets. To standardise the universal dataset, only markers which are present in the output of the Wald test (genes with variance greater than zero among samples) were used. Similarly, to enable fair comparisons, only differentially expressed protein coding genes and documented lncRNAs were used from the DEG lists, as was surveyed in the cell type specific marker paper. Bonferroni correction was applied to the hypergeometric distribution p values to account for multiple testing and significance scores were calculated using -log10(corrected p value). For the mapping of marker genes to the interaction networks, no filters were applied.

### Reconstruction of molecular networks

Mice regulatory networks containing directed regulatory layers were retrieved from multiple databases (Table S6): miRNA-mRNA (ie., miRNAs regulating mRNAs) and lncRNA-miRNA (ie., lncRNAs regulating miRNAs) interactions were downloaded from TarBase v7.0 (113) and LncBase v2.0 (114), respectively. Only miRNA-mRNA and lncRNA-miRNA interactions determined using HITS-CLIP (115) experiments were considered. Regulatory interactions between transcription factors (TFs) and miRNAs (ie. TFs regulating miRNAs) were retrieved from TransmiR v1.2 (116), GTRD (117) and TRRUST v2 (118,119). Co-expression based inferences were ignored from all the above resources. Transcriptional regulatory interactions (ie., TFs regulating target genes) were inferred using data from ORegAnno 3.0 (61), GTRD and TRUSST. In cases where transcriptional regulatory interactions are derived from high-throughput datasets such as ChIP-seq, we attributed the regulatory interaction elicited by the bound transcription factor to genes which lie within a 10kb window on either side of the ChIP-seq peak (ORegAnno) or meta-cluster (in the case of GTRD). TF-lncRNA interactions (ie., TFs regulating lncRNAs) were also inferred based on the ChIP-seq binding profiles represented by meta-clusters in GTRD. We used only TF-lncRNA interactions within intergenic lncRNAs to avoid assigning false regulatory interactions due to the high number of instances where the lncRNAs overlap with protein-coding genes. In addition, no overlaps were allowed between the coordinates of the ChIP-seq peaks / meta-clusters and any gene annotation. Only if the first annotation feature within a 10kb genomic window downstream to the ChIP-seq peak / meta-cluster was designated as an intergenic lncRNA, a regulatory interaction between the TF and the lncRNA was assigned. Bedtools (120) was used for the custom analyses to look for overlaps between coordinates. All the nodes in the collected interactions were represented by their Ensembl gene IDs for standardization. A summary of the interactions collected from each resource and the quality control criteria applied is given in Table S6.

To generate PCeE and GCeE regulatory networks, interactions in this collated universal network were filtered using the differential expression data (Figure 1). The assumption was made that if both nodes of a particular interaction were expressed in the RNAseq data, the interaction is possible. Furthermore, to filter for the interactions of prime interest, only nodes which were differentially expressed and their associated interactors were included in the regulatory networks.

### Cluster analysis

Clusters of highly interconnected regions within the PCeE and GCeE regulatory networks were identified using the MCODE plugin within Cytoscape (37,121). Default settings were applied: degree cutoff = 2, haircut = true, node score cutoff = 0.2, k-core = 2 and max depth = 100. Clusters were visualised in Cytoscape.

The nodes of each cluster were tested for functional enrichment (hypergeometric model, q value ≤ 0.05) based on Reactome annotations using the ReactomePA R package (38,107–109) following conversion from mouse to human identifiers using Inparanoid (v8) (110,111). Cases where the number of nodes associated with a pathway < 5 were considered not significant regardless of the q value. The top 5 significant Reactome pathways associated with each cluster were visualised using a heatplot generated in R (Figure 3B). More than 5 pathways were visualised, where multiple Reactome pathways had equal q values.

### Master regulator analysis

To identify potential master regulators of the Paneth cell and the goblet cell types, the upstream regulators of cell type specific markers (from (12)) were investigated. To do this, all markers were mapped to the relevant networks then subnetworks were extracted consisting of markers and their regulators (Table S7).

### Regulatory rewiring analysis

To calculate rewiring scores for regulators, sub-networks were extracted (from the PCeE and GCeE regulatory networks) containing just the regulator of interest and its downstream targets. For each regulator of interest, the subnetworks from the PCeE and GCeE networks were compared using the Cytoscape app DyNet (121,122). The degree corrected Dn score was extracted for each regulator and used to quantify rewiring of the regulator’s downstream targets between the PCeE and GCeE regulatory networks. Functional analysis was carried out on the targets of the top five most rewired regulators. For each regulator, the targets were classified based on whether they are present in only the PCeE network, only the GCeE network or in both networks. Each group of targets was tested for functional enrichment (hypergeometric model, q value ≤ 0.1) based on Reactome and KEGG annotations using the ReactomePA R package (38,107–109) following conversion from mouse to human identifiers using Inparanoid (v8) (110,111).

### IBD and drug target associated genes

Genes associated with UC and CD based on single nucleotide polymorphisms were obtained from two studies (67,68). Additionally, the top 100 differentially expressed genes were obtained from goblet cell analysis of inflamed UC vs healthy human colonic tissue from (13). Genes were converted to Mouse Ensembl identifiers using Inparanoid (v8) and bioDBnet db2db (110–112). Additionally, to enable hypergeometric significant testing with the universal network as the background, only UC and CD genes present in the universal network are included in the analyses. eQTL datasets for CD were retrieved from (69) while the list of targets related to drug-interactions was downloaded from (123).

### Quantification and statistical analysis

Statistical parameters including the exact value of n and statistical significance are reported in the Figures and Figure Legends. n represents the number of enteroid biological replicates generated. Where relevant, data is judged to be statistically significant when Bonferroni corrected p value ≤ 0.01. Genes with absolute log2 fold change of ≥ |1| and false discovery rate ≤ 0.05 were considered to be differentially expressed. Based on principal component analysis of transcript expression, one biological replicate from the Paneth cell enriched enteroids was identified as an outlier and removed (Figure S3). Where stated, the hypergeometric distribution model was used to calculate significance using R.

### Data and software availability

Small and stranded RNA-seq data has been deposited in the European Nucleotide Archive (ENA) with accession numbers PRJEB32354 and PRJEB32366 respectively. Scripts to analyse the differentially expressed genes are available on GitHub: https://github.com/korcsmarosgroup/organoid_regulatory_networks.

## Supplementary items

**Table S1: Differentially expressed genes from cell type enriched enteroids vs conventionally differentiated organoids** (q value ≤ 0.05 and log2 fold change ≥ |1|). Gene annotations included.

**Table S2: Expression and differential expression values for primary cell type markers.** Expression given as mean transcripts per million (TPM) for each enteroid type. Log2 fold changes values given where differential expression criteria passed (q value ≤ 0.05 and log2 fold change ≥ |1|). Control-normally differentiated enteroids; Paneth-Paneth cell enriched enteroids; goblet-goblet cell enriched enteroids.

**Table S3: Differentially expressed antimicrobial peptide (AMP) and mucin related genes in Paneth cell enriched enteroids and goblet cell enriched enteroids (compared to conventionally differentiated enteroids).** Only genes which are differentially expressed (log2fc ≥ 1 and false discovery rate ≤ 0.05) in at least one of the datasets was included. Lfc = log2 fold change; fdr = false discovery rate; DEG = differentially expressed gene; Paneth = Paneth enriched enteroid, goblet = goblet enriched enteroid.

**Table S4: Hypergeometric distribution testing of cell type specific marker enrichment in upregulated differentially expressed gene lists.**

**Table S5: Functional enrichment analysis of the PCeE and GCeE clusters (q val** ≤ 0.1).

**Table S6: A summary of the molecular interactions compiled to generate the universal network.**

**Table S7: Interactions between markers and their regulators in the Paneth cell and the goblet cell subnetworks, including regulator specificity.**

**Table S8: Rewiring analysis results for the marker regulators present in the Paneth cell and the goblet cell subnetworks.** Dn score generated using Cytoscape app DyNet.

**Table S9: Functional enrichment analysis of the top five most rewired (shared) marker regulators (q val ≤ 0.1).**

**Table S10: Crohn’s disease SNP associated genes in the enriched enteroid regulatory networks.**

**Table S11: Ulcerative colitis SNP associated genes in the enriched enteroid regulatory networks.**

**Table S12: Crohn’s disease eQTL associated genes and drug target genes in the enriched enteroid regulatory networks.**

**Table S13: IBD associated genes targeted by predicted master regulators in the enriched enteroid regulatory networks.** Ulcerative colitis (UC) and Crohn’s disease (CD) associated genes (from SNP data) targeted by at least one of the master regulators in the relevant networks; list of top 100 CD differentially expressed genes in human colonic biopsies (CD inflamed vs healthy) which are targeted by at least one of the predicted goblet cell master regulators in the GCeE network.

**Figure S1: Small intestinal 3D organoid culture. A.** Culture of isolated mouse small intestinal epithelial crypts in Matrigel matrix and ENR media (conventionally differentiated) for 7 days. Isolated crypts form 3D cysts which bud after 2 days of culture to form crypt- and villus-like domains. Paneth cells are clearly visible by light microscopy (Black arrows). Mucous and shedding cells accumulate in the central lumen of organoids (*). n = 3. **B.** cell type specific enrichment illustrated by immunofluorescence labelling of cultured mouse 3D enteroids, conventionally differentiated (left) and enriched for either Paneth cells or goblet cells (right). Lysozyme granules characteristic of Paneth cells are indicated with a green arrow. Goblet cells were identified using a specific anti-Muc2 mucin antibody (pink).

**Figure S2: Transcript abundances and differential expression of five major cell type markers. A:** Mean transcript abundances in the conventionally differentiated, goblet cell enriched and Paneth cell enriched enteroids. B: Log2 fold change in the goblet cell enriched enteroid vs conventional enteroid analysis and the Paneth cell enriched enteroid vs conventional enteroid analysis. Data only presented where the differential expression criteria passed (q value ≤ 0.05 and log2 fold change ≥|1|).

**Figure S3: Principal component analysis of Paneth cell enriched enteroid transcriptomics data from each biological replicate.**

**Item S1: Cytoscape file for the Paneth cell enriched enteroid regulatory network, including sub-clusters.**

**Item S2: Cytoscape file for the goblet cell enriched enteroid regulatory networks, including sub-clusters.**

