## Supplementary material for "Regulatory network analysis of Paneth cell and goblet cell enriched gut organoids using transcriptomics approaches": Tables S2-4, 6, 8, 10, 11, 13, Figures S1-3

Supplementary Items

| Marker gene | Cell type specificity | Mean expression (TPM) | | | Log2 fold change | |
| --- | --- | --- | --- | --- | --- | --- |
|  |  | control | goblet | Paneth | goblet v ctrl | Paneth v ctrl |
| *Lgr5* | Stem | 71.88 | 4.08 | 12.76 | -2.77 | -1.16 |
| *ChgA* | Enteroendocrine | 56.49 | 507.58 | 287.2 | 2.42 | 2.17 |
| *Muc2* | Goblet | 47.26 | 625.05 | 185.18 | 2.81 | 1.83 |
| *Cd24a* | Paneth | 471.23 | 1034.85 | 708.78 | 1.05 | NA |
| *Lyz1* | Paneth | 2613.91 | 11576.37 | 22065.95 | 1.72 | 2.67 |
| *Vil1* | Epithelial | 211.13 | 219.22 | 89.02 | NA | NA |

**Table S2:** **Expression and differential expression values for primary cell-type markers.** Expression given as mean transcripts per million (TPM) for each enteroid type. Log2 fold changes values given where differential expression criteria passed (q value ≤ 0.05 and log2 fold change ≥ |1|). Control- normally differentiated enteroids; Paneth= Paneth cell enriched enteroids; goblet= goblet cell enriched enteroids.

| **Gene Type** | **Gene Name** | **Ensembl ID** | **Paneth DEG lfc** | **Paneth DEG fdr** | **Goblet DEG lfc** | **Goblet DEG fdr** |
| --- | --- | --- | --- | --- | --- | --- |
| AMP | *Ang4* | ENSMUSG00000060615 | 4.64 | 8.61E-36 | 2.45 | 8.75E-13 |
| AMP | *Defa17* | ENSMUSG00000060208 | 4.02 | 6.40E-46 | 2.95 | 5.56E-31 |
| AMP | *Defa2* | ENSMUSG00000096295 | 4.58 | 1.70E-18 | NA | NA |
| AMP | *Defa20* | ENSMUSG00000095066 | 4.40 | 1.19E-17 | NA | NA |
| AMP | *Defa21* | ENSMUSG00000074447 | 5.18 | 2.22E-22 | NA | NA |
| AMP | *Defa22* | ENSMUSG00000074443 | 5.40 | 3.09E-20 | NA | NA |
| AMP | *Defa23* | ENSMUSG00000074446 | 3.69 | 1.94E-12 | NA | NA |
| AMP | *Defa24* | ENSMUSG00000064213 | 3.91 | 8.51E-38 | 3.24 | 1.92E-32 |
| AMP | *Defa26* | ENSMUSG00000060070 | 3.02 | 1.86E-39 | 2.22 | 8.61E-27 |
| AMP | *Defa28* | ENSMUSG00000074434 | 2.84 | 3.21E-17 | 1.66 | 1.20E-07 |
| AMP | *Defa29* | ENSMUSG00000074437 | 1.89 | 3.20E-06 | NA | NA |
| AMP | *Defa3* | ENSMUSG00000074440 | 4.01 | 6.69E-26 | 2.90 | 4.28E-17 |
| AMP | *Defa30* | ENSMUSG00000074444 | 3.89 | 1.94E-21 | 1.44 | 3.15E-4 |
| AMP | *Defa32* | ENSMUSG00000094818 | 5.54 | 1.06E-13 | NA | NA |
| AMP | *Defa33* | ENSMUSG00000094362 | 5.35 | 9.86E-13 | NA | NA |
| AMP | *Defa34* | ENSMUSG00000063206 | 5.34 | 5.97E-57 | 2.32 | 1.00E-13 |
| AMP | *Defa35* | ENSMUSG00000061845 | 5.85 | 3.04E-20 | 1.43 | 0.03 |
| AMP | *Defa36* | ENSMUSG00000094662 | 4.52 | 7.61E-37 | 2.16 | 7.04E-11 |
| AMP | *Defa5* | ENSMUSG00000074439 | 4.97 | 7.91E-33 | NA | NA |
| AMP | *Lyz1* | ENSMUSG00000069515 | 3.86 | 6.73E-27 | 2.49 | 3.67E-14 |
| AMP | *Pla2g2a* | ENSMUSG00000058908 | 3.23 | 6.78E-45 | NA | NA |
| AMP | *Reg3g* | ENSMUSG00000074447 | 5.18 | 2.22E-22 | NA | NA |
| Mucin related | *Fcgbp* | ENSMUSG00000047730 | 2.02 | 5.22E-08 | 4.40 | 8.45E-44 |
| Mucin related | *Muc1* | ENSMUSG00000042784 | NA | NA | NA | NA |
| Mucin related | *Muc13* | ENSMUSG00000022824 | NA | NA | 1.03 | 2.48E-4 |
| Mucin related | *Muc2* | ENSMUSG00000025515 | 2.64 | 8.57E-10 | 4.06 | 2.24E-27 |
| Mucin related | *Muc3* | ENSMUSG00000037390 | -2.35 | 0.02 | NA | NA |
| Mucin related | *Muc3a* | ENSMUSG00000094840 | 1.59 | 3.46E-06 | 2.46 | 1.09E-16 |
| Mucin related | *Retnlb* | ENSMUSG00000022650 | NA | NA | NA | NA |
| Mucin related | *Tff3* | ENSMUSG00000024029 | 3.27 | 1.89E-26 | 3.69 | 6.74E-42 |

**Table S3: Differentially expressed antimicrobial peptide (AMP) and mucin related genes in Paneth cell enriched enteroids and goblet cell enriched enteroids (compared to conventionally differentiated enteroids).** Only genes which are differentially expressed (log2fc ≥ 1 and false discovery rate ≤ 0.05) in at least one of the datasets was included. Lfc = log2 fold change; fdr = false discovery rate; DEG = differentially expressed gene; Paneth = Paneth enriched enteroid, goblet = goblet enriched enteroid.

| **Marker list cell type** | **DEG list** | **#Markers** | **#DEGs lncRNAs & protein coding** | **#DEG Markers** | **Hypergeometric pval** | **Multiple testing corrected Pan&Gob** | **Multiple testing corrected all cell types** | **-Log10(qval) Pan&Gob** | **-Log10(qval) all** |
| --- | --- | --- | --- | --- | --- | --- | --- | --- | --- |
| Paneth | Paneth | 71 | 2077 | 56 | 6.07E-36 | 2.43E-35 | 3.64E-35 | 34.615 | 34.438 |
| goblet | Paneth | 422 | 2077 | 102 | 7.69E-10 | 3.07E-09 | 4.61E-09 | 8.512 | 8.336 |
| enteroendocrine | Paneth | 204 | 2077 | 140 | 7.32E-75 | NA | 4.39E-74 | NA | 73.357 |
| Paneth | Goblet | 71 | 1797 | 40 | 6.97E-20 | 2.79E-19 | 4.18E-19 | 18.555 | 18.379 |
| goblet | Goblet | 422 | 1797 | 173 | 1.01E-55 | 4.05E-55 | 6.08E-55 | 54.392 | 54.216 |
| enteroendocrine | Goblet | 204 | 1797 | 148 | 8.14E-94 | NA | 4.89E-93 | NA | 92.311 |

**Table S4:** **Hypergeometric significance testing of cell type specific marker enrichment in upregulated differentially expressed gene lists.**

| **Interaction type** | **Source(s)** | **Number of unique interactions** | **Quality control criteria** |
| --- | --- | --- | --- |
| TF-TG | TRRUST v2  GTRD  ORegAnno v3.0 | 1066383 | - ChIP-Seq peaks should not overlap any gene annotation; if peak on + strand, only the first gene downstream to the gene or if peak on - strand, only first gene upstream to the peak is considered. - Genes attributed to the transcription factor which lie within a 10kb window on either side of the ChIP-seq peak (ORegAnno) or meta-cluster (in the case of GTRD). |
| TF-lncRNA | GTRD | 159055 | - ChIP-Seq peaks should not overlap any gene annotation; if peak on + strand, only the first gene downstream to the gene or if peak on - strand, only first gene upstream to the peak is considered. - Genes attributed to the transcription factor which lie within a 10kb window on either side of the meta-cluster. - Only if the first annotation feature within a 10kb genomic window downstream to the ChIP-seq peak / meta-cluster was designated as an intergenic lncRNA, a regulatory interaction between the TF and the lncRNA was assigned - to avoid assigning false regulatory interactions due to the high number of instances where the lncRNAs overlap with protein-coding genes. |
| miRNA-mRNA | TarBase v7.0 | 141892 | - Only HITS-CLIP based experimental evidence considered. - Co-expression based inferences not considered. |
| TF-miRNA | TransmiR v1.2  TRRUST v2  GTRD | 9204 | - ChIP-Seq peaks should not overlap any gene annotation; if peak on + strand, only the first gene downstream to the gene or if peak on - strand, only first gene upstream to the peak is considered. - Co-expression based inferences not considered |
| lncRNA - miRNA | lncBase2 | 6637 | - Only HITS-CLIP based experimental evidence considered. - Co-expression based inferences not considered. |

**Table S6:** **A summary of the physical interactions compiled to generate the universal network.**

| **Shared regulator** | **Shared regulator_id** | **D_n_-Score (degree corrected)** | **# Shared targets** | **# Pan only targets** | **# Gob only targets** |
| --- | --- | --- | --- | --- | --- |
| Etv4 | ENSMUSG00000017724 | 0.4 | 1 | 3 | 1 |
| mmu-let-7e-5p | mmu-let-7e-5p | 0.37037037 | 49 | 102 | 38 |
| mmu-miR-152-3p | mmu-miR-152-3p | 0.352112676 | 63 | 104 | 46 |
| Myb | ENSMUSG00000019982 | 0.339147287 | 83 | 108 | 67 |
| Rora | ENSMUSG00000032238 | 0.330926594 | 281 | 386 | 164 |
| Mitf | ENSMUSG00000035158 | 0.330246914 | not calculated | not calculated | not calculated |
| Hoxb4 | ENSMUSG00000038692 | 0.314465409 | not calculated | not calculated | not calculated |
| Nr5a2 | ENSMUSG00000026398 | 0.317622951 | not calculated | not calculated | not calculated |
| Irf1 | ENSMUSG00000018899 | 0.317073171 | not calculated | not calculated | not calculated |
| mmu-miR-7a-5p | mmu-miR-7a-5p | 0.3125 | not calculated | not calculated | not calculated |
| Foxa1 | ENSMUSG00000035451 | 0.311414392 | not calculated | not calculated | not calculated |
| Tead4 | ENSMUSG00000030353 | 0.3097313 | not calculated | not calculated | not calculated |
| Nkx2-2 | ENSMUSG00000027434 | 0.306954436 | not calculated | not calculated | not calculated |
| Vdr | ENSMUSG00000022479 | 0.302962662 | not calculated | not calculated | not calculated |
| Ets1 | ENSMUSG00000032035 | 0.302570586 | not calculated | not calculated | not calculated |
| Nr3c1 | ENSMUSG00000024431 | 0.302290333 | not calculated | not calculated | not calculated |
| Foxa3 | ENSMUSG00000040891 | 0.301075269 | not calculated | not calculated | not calculated |
| Bhlha15 | ENSMUSG00000052271 | 0.297066015 | not calculated | not calculated | not calculated |
| mmu-miR-101a-3p | mmu-miR-101a-3p | 0.290789474 | not calculated | not calculated | not calculated |
| Zfp57 | ENSMUSG00000036036 | 0.28757764 | not calculated | not calculated | not calculated |
| Fosl1 | ENSMUSG00000024912 | 0.292517007 | not calculated | not calculated | not calculated |
| Pax6 | ENSMUSG00000027168 | 0.288018433 | not calculated | not calculated | not calculated |
| Nfatc2 | ENSMUSG00000027544 | 0.295454545 | not calculated | not calculated | not calculated |
| Neurod1 | ENSMUSG00000034701 | 0.280855199 | not calculated | not calculated | not calculated |
| Insm1 | ENSMUSG00000068154 | 0.281121751 | not calculated | not calculated | not calculated |
| mmu-miR-153-3p | mmu-miR-153-3p | 0.272727273 | not calculated | not calculated | not calculated |
| Neurod2 | ENSMUSG00000038255 | 0.269685039 | not calculated | not calculated | not calculated |
| Fosb | ENSMUSG00000003545 | 0.265217391 | not calculated | not calculated | not calculated |
| Klf15 | ENSMUSG00000030087 | 0.285714286 | not calculated | not calculated | not calculated |
| Atoh1 | ENSMUSG00000073043 | 0.244949495 | not calculated | not calculated | not calculated |

**Table S8: Rewiring analysis results for the marker regulators present in the Paneth and the goblet subnetworks.** D_n_ score generated using Cytoscape app DyNet.

| **Cell-type specific regulatory network** | **Crohn's susceptibility gene** | **Direction of differential expression** |
| --- | --- | --- |
| Paneth | 9430076C15Rik | Upregulated |
| Paneth | Atg16l2 | Upregulated |
| Paneth | Fut2 | Upregulated |
| Paneth | Hmha1 | Upregulated |
| Paneth | Itln1 | Upregulated |
| Paneth | Izumo1 | Upregulated |
| Paneth | Jazf1 | Upregulated |
| Paneth | Plcl1 | Upregulated |
| Paneth | Tnfsf15 | Upregulated |
| Paneth | Ccdc88b | Downregulated |
| Paneth | Dbp | Downregulated |
| Paneth | Fads1 | Downregulated |
| Paneth | Fads2 | Downregulated |
| Paneth | H2-Q1 | Downregulated |
| Paneth | H2-Q10 | Downregulated |
| Paneth | H2-Q2 | Downregulated |
| Paneth | H2-Q6 | Downregulated |
| Paneth | H2-Q7 | Downregulated |
| Paneth | Kif21b | Downregulated |
| Paneth | Ksr1 | Downregulated |
| Paneth | Ptpn22 | Downregulated |
| Paneth | Zpbp2 | Downregulated |
| Goblet | Fut2 | Upregulated |
| Goblet | Hmha1 | Upregulated |
| Goblet | Inpp5d | Upregulated |
| Goblet | Itln1 | Upregulated |
| Goblet | Izumo1 | Upregulated |
| Goblet | Jazf1 | Upregulated |
| Goblet | Plcl1 | Upregulated |
| Goblet | Tnfsf15 | Upregulated |
| Goblet | Gart | Downregulated |
| Goblet | H2-Q7 | Downregulated |
| Goblet | H2-Q6 | Downregulated |
| Goblet | Notch2 | Downregulated |

**Table S10: Crohn’s disease SNP associated genes in the enriched enteroid regulatory networks.**

| **Cell-type specific regulatory network** | **Ulcerative Colitis susceptibility gene** | **Direction of differential expression** |
| --- | --- | --- |
| Paneth | Dap | Upregulated |
| Paneth | Edem2 | Upregulated |
| Paneth | Itgal | Upregulated |
| Paneth | Maml2 | Upregulated |
| Paneth | Mmp24 | Upregulated |
| Paneth | Nr5a2 | Downregulated |
| Paneth | Plcl1 | Upregulated |
| Paneth | Tnfsf15 | Upregulated |
| Paneth | Zpbp2 | Downregulated |
| Paneth | Card11 | Downregulated |
| Paneth | Hnf4A | Downregulated |
| Paneth | Nusap1 | Downregulated |
| Paneth | Procr | Upregulated |
| Goblet | Dap | Upregulated |
| Goblet | Edem2 | Upregulated |
| Goblet | Itgal | Upregulated |
| Goblet | Mmp24 | Upregulated |
| Goblet | Nr5a2 | Downregulated |
| Goblet | Plcl1 | Upregulated |
| Goblet | Tnfsf15 | Upregulated |
| Goblet | Card11 | Downregulated |
| Goblet | Cep250 | Downregulated |
| Goblet | Procr | Upregulated |

**Table S11: Ulcerative colitis disease SNP associated genes in the enriched enteroid regulatory networks.**

| **UC SNP associated genes** | **CD SNP associated genes** | **Goblet differentially expressed genes** |
| --- | --- | --- |
| ENSMUSG00000026398 | ENSMUSG00000053007 | ENSMUSG00000013523 |
| ENSMUSG00000027611 | ENSMUSG00000010663 | ENSMUSG00000017057 |
| ENSMUSG00000027612 | ENSMUSG00000017195 | ENSMUSG00000024597 |
| ENSMUSG00000030830 | ENSMUSG00000018334 | ENSMUSG00000027006 |
| ENSMUSG00000036526 | ENSMUSG00000024665 | ENSMUSG00000027346 |
| ENSMUSG00000038241 | ENSMUSG00000027843 | ENSMUSG00000027513 |
| ENSMUSG00000038349 | ENSMUSG00000038349 | ENSMUSG00000027876 |
| ENSMUSG00000039168 | ENSMUSG00000047767 | ENSMUSG00000028236 |
| ENSMUSG00000050395 | ENSMUSG00000047810 | ENSMUSG00000031844 |
| ENSMUSG00000038312 | ENSMUSG00000050395 | ENSMUSG00000032322 |
|  | ENSMUSG00000055978 | ENSMUSG00000032978 |
|  | ENSMUSG00000059824 | ENSMUSG00000034472 |
|  | ENSMUSG00000060550 | ENSMUSG00000038039 |
|  | ENSMUSG00000063568 | ENSMUSG00000039234 |
|  | ENSMUSG00000067235 | ENSMUSG00000046841 |
|  | ENSMUSG00000073409 | ENSMUSG00000055976 |
|  | ENSMUSG00000079507 | ENSMUSG00000074004 |
|  | ENSMUSG00000038209 | ENSMUSG00000075610 |
|  | ENSMUSG00000091705 | ENSMUSG00000055963 |
|  | ENSMUSG00000035697 | ENSMUSG00000036764 |
|  | ENSMUSG00000064158 |  |

**Table S13: IBD associated genes targeted by predicted master regulators in the enriched enteroid regulatory networks.** Ulcerative colitis (UC) and Crohn’s disease (CD) associated genes (from SNP data) targeted by at least one of the master regulators in the relevant networks; list of top 100 CD differentially expressed genes in human colonic biopsies (CD inflamed vs healthy) which are targeted by at least one of the predicted goblet cell master regulators in the GCeE network.

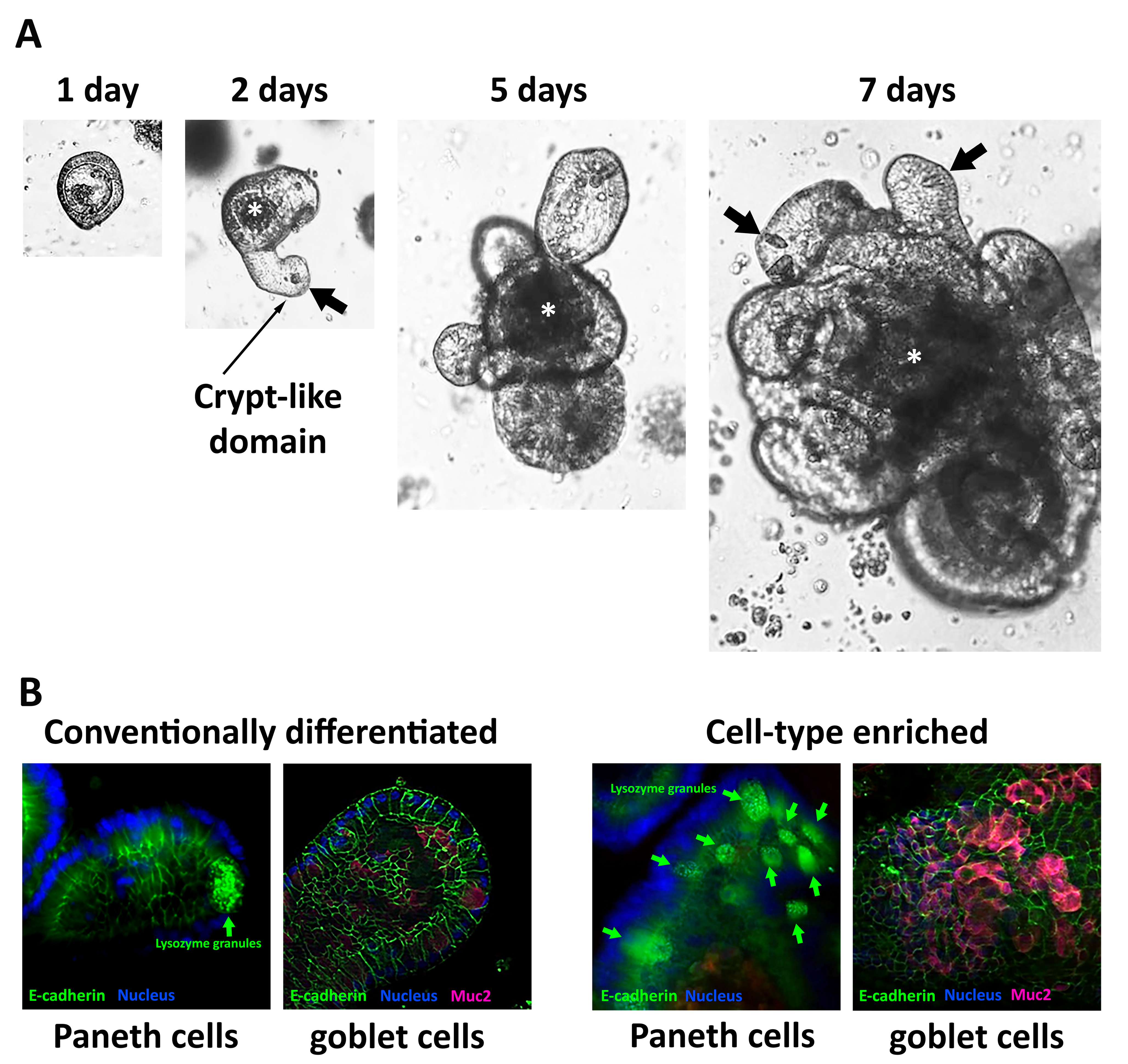

**Figure S1: Small intestinal 3D organoid culture.** **A.** Culture of isolated mouse small intestinal epithelial crypts in Matrigel matrix and ENR media (conventionally differentiated) for 7 days. Isolated crypts form 3D cysts which bud after 2 days of culture to form crypt- and villus-like domains. Paneth cells are clearly visible by light microscopy (Black arrows). Mucous and shedding cells accumulate in the central lumen of organoids (*). n = 3. **B.** cell type specific enrichment illustrated by immunofluorescence labelling of cultured mouse 3D enteroids, conventionally differentiated (left) and enriched for either Paneth cells or goblet cells (right). Lysozyme granules characteristic of Paneth cells are indicated with a green arrow. Goblet cells were identified using a specific anti-Muc2 mucin antibody (pink).

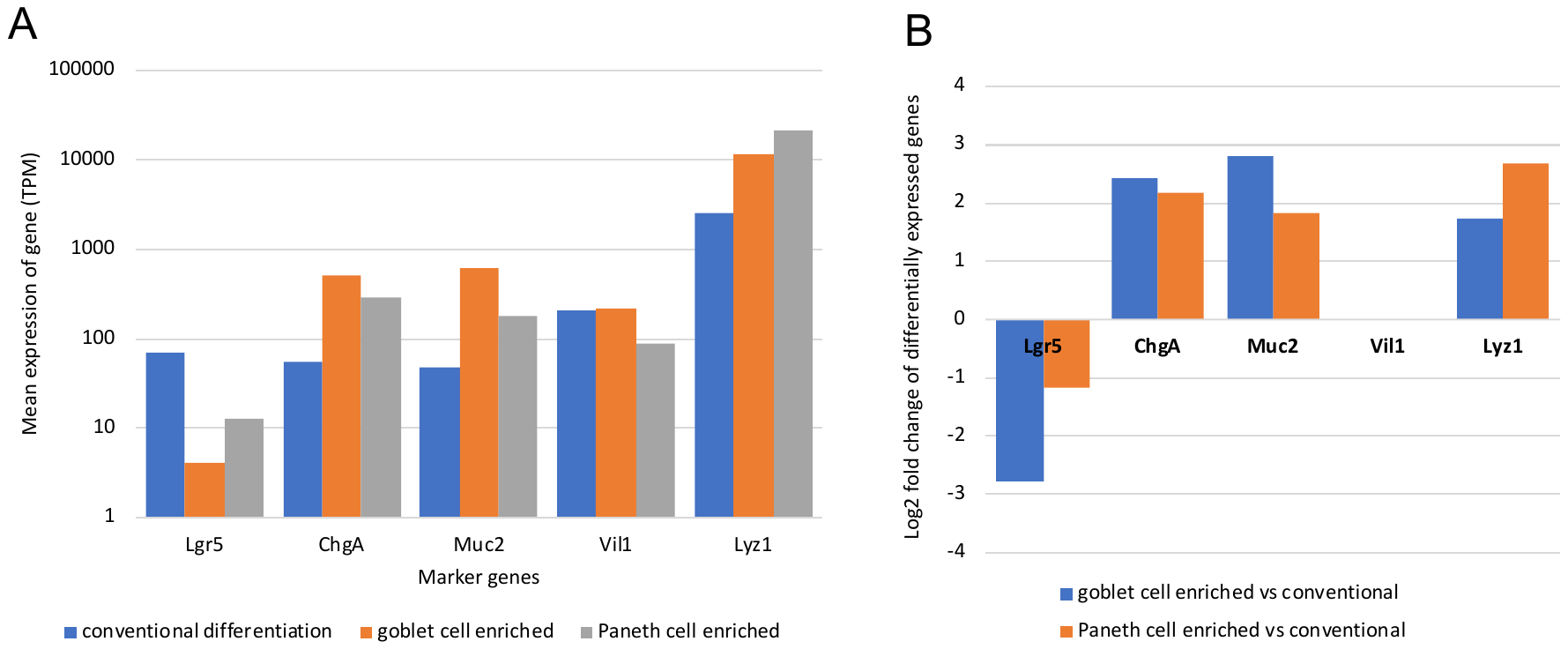

**Figure S2:** **Transcript abundances and differential expression of five major cell-type markers.** **A:** Mean transcript abundances in the conventionally differentiated, goblet cell enriched and Paneth cell enriched enteroids. **B:** Log2 fold change in the goblet cell enriched enteroid vs conventional enteroid analysis and the Paneth cell enriched enteroid vs conventional enteroid analysis. Data only presented where the differential expression criteria passed (q value ≤ 0.05 and log2 fold change ≥|1|).

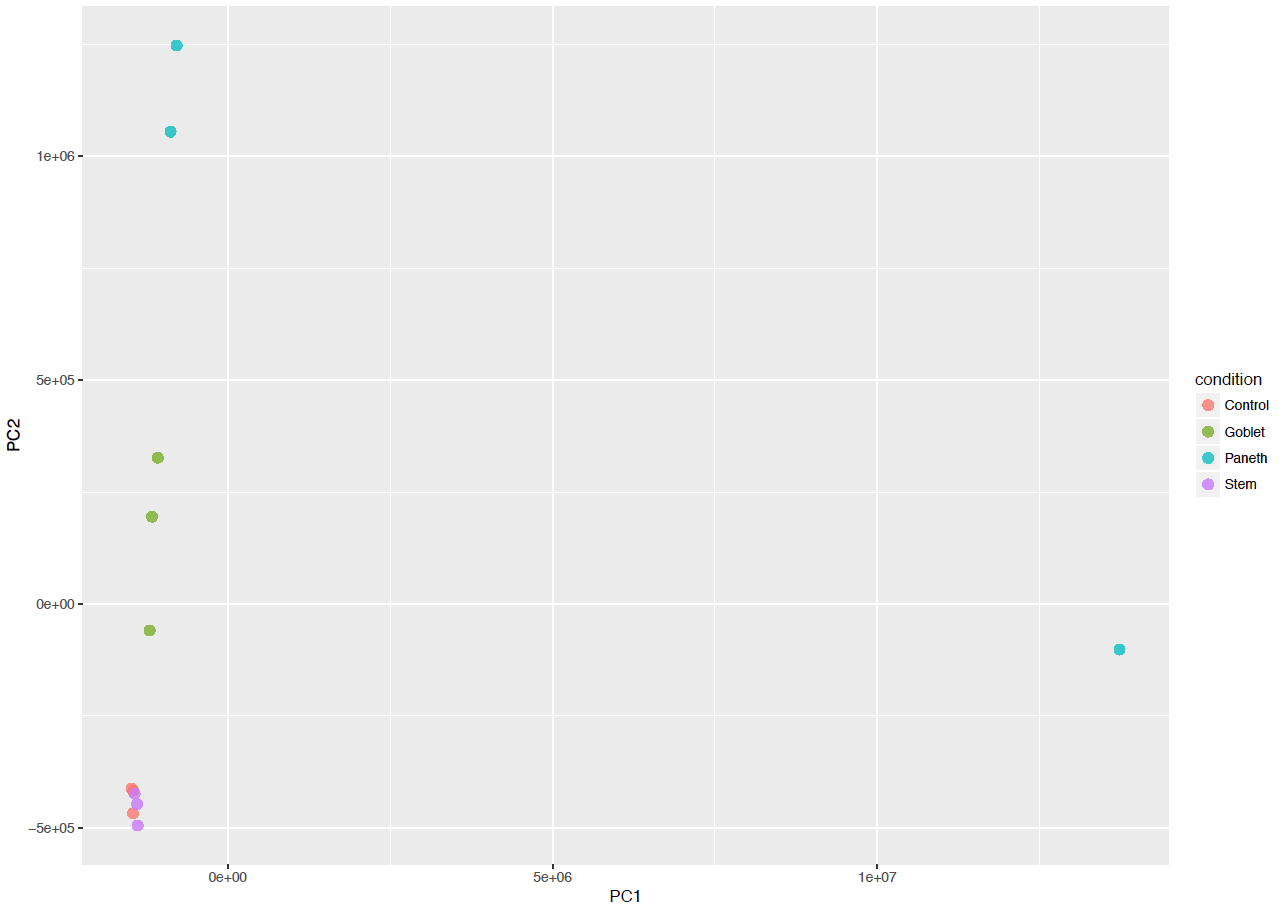

**Figure S3: Principal component analysis of Paneth cell enriched enteroid transcriptomics data from each biological replicate.**
